# The Ribb-osome: Ribbon boosts ribosomal protein gene expression to coordinate organ form and function

**DOI:** 10.1101/2021.10.27.466115

**Authors:** Rajprasad Loganathan, Daniel C. Levings, Ji Hoon Kim, Michael B. Wells, Hannah Chiu, Yifan Wu, Matthew Slattery, Deborah J. Andrew

## Abstract

Cell growth is well defined for the late (post-embryonic) stages of development, but evidence for early (embryonic) cell growth during post-mitotic morphogenesis is quite limited. Here, we identify early cell growth as a key characteristic of tubulogenesis in the *Drosophila* embryonic salivary gland (SG). A BTB/POZ domain nuclear factor, Ribbon (Rib), mediates this early cell growth. Rib binds the transcription start site of nearly every SG-expressed ribosomal protein gene (RPG) and is required for full expression of all RPGs tested. Rib binding to RPG promoters *in vitro* is weak and not sequence-specific, suggesting that specificity is achieved through co-factor interactions. Consistent with this hypothesis, we demonstrate Rib’s ability to physically interact with each of the three known contributors to RPG transcription. Surprisingly, Rib-dependent early cell growth in another tubular organ—the embryonic trachea—is not mediated by direct RPG transcription. These findings support a model of early cell growth sustained by transcriptional regulatory networks customized for organ form and function.

## INTRODUCTION

Epithelial tubes are vital to metazoan physiological processes, *e.g.*, fluid secretion, storage, absorption, exchange, and transport. Epithelial tubes originate from all three germ layers: ectoderm (*e.g.* mammary gland), endoderm (*e.g.* lung), and mesoderm (*e.g.* kidney); and have both branched (*e.g.* vasculature) and unbranched (*e.g.* gut) architectures. Studies conducted across the metazoan phyla in several model systems have uncovered shared molecular and cellular mechanisms of tubulogenesis, *e.g.* fibroblast growth factor-dependent branching morphogenesis (Danopoulos et al., 2019; Klambt et al., 1992; Ohshiro et al., 2002; Reichman-Fried et al., 1994; Reichman-Fried and Shilo, 1995; Sutherland et al., 1996; Walker et al., 2016). Previous studies have also revealed mechanisms of tube assembly specifically adapted to organ functional requirements, *e.g.*, integrin-dependent reversal of junctional polarity determinants in the *Drosophila* midgut to facilitate absorption and to maintain epithelial integrity in the absence of a protective luminal matrix (Chen et al., 2018). Morphogenetic programs may, hence, utilize both shared and tissue-specific mechanisms in making and shaping epithelial tubes.

Of the various model systems of epithelial tubulogenesis, few exhibit the advantages of both genetic manipulability and relative simplicity of access for morphological and molecular-level characterizations as the *Drosophila* embryonic salivary gland (SG) (Chung et al., 2014; Sidor and Röper, 2016). SG morphogenesis is post-mitotic and non-apoptotic (Campos-Ortega and Hartenstein, 1997; Poulson, 1937, 1950), rendering its construction process exemplary for investigations of molecular and cellular mechanisms of epithelial tubulogenesis exclusively by changes in cell position, shape, or size. SGs are first visible during embryonic stage 10 as two epithelial placodes on either side of the midline on the ventral surface of parasegment 2 (**Fig 1A).** SG assembly begins during embryonic stage 11, with the sequential invagination of cells to form a nascent tube (Booth et al., 2014; Chung et al., 2017; Myat and Andrew, 2000b). Tube maturation is marked by elongation via oriented apical membrane expansion and cell rearrangement (Sanchez-Corrales et al., 2018; Xu et al., 2011), with concomitant organ positioning by integrin-dependent collective cell migration beginning at stage 12 and continuing through stage 16 (Bradley et al., 2003). Upon maturation, each of the pair of SGs contains an unbranched secretory tube, which connects to a duct tube at the proximal end (**Fig. 1B**).

**Figure 1.**
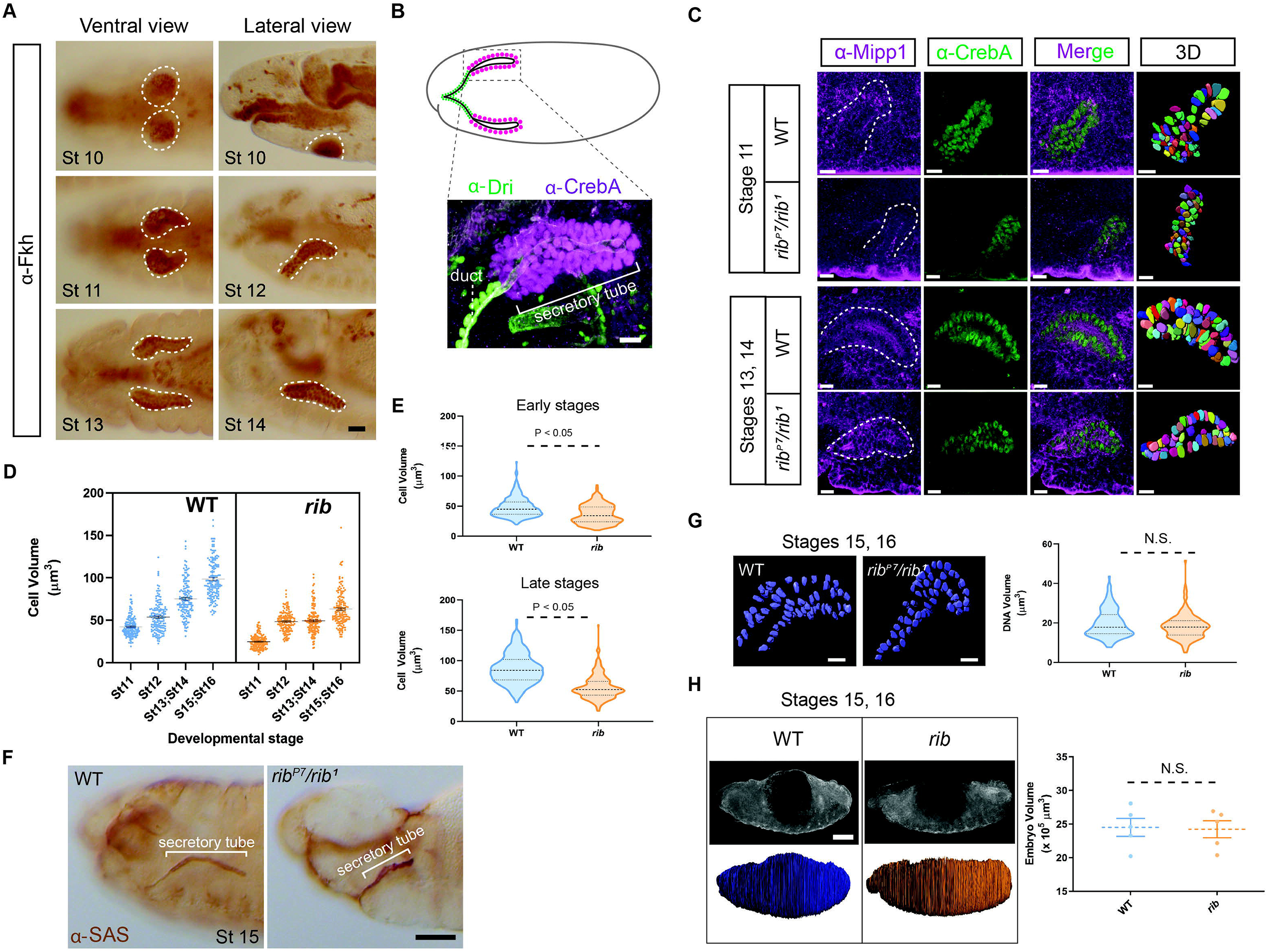
Rib is required for cell growth in the embryonic salivary gland (SG). **(A)** SG (outline) tube morphology at distinct embryonic stages. Placode cells (stage 10) internalize to form an incipient tube (stage 11). Elongation and posterior turning (stage 12) positions the tube along the anteroposterior axis as the cells collectively migrate during tube maturation (stages 13, 14). Scale bar: 10 µm. **(B)** The bilateral secretory tubes (magenta nuclei) of the late embryonic SG connect with each other and to the digestive tract via the ducts (green nuclei). Scale bar: 5 µm. **(C)** 3D rendering and SG cell volumetry of embryos stained with α-Mipp1 (magenta, cell boundaries) and α-CrebA (green, nuclei) in WT and *rib* mutant SGs. Scale bars: 10 µm. **(D)** SG cell size is reduced in *rib* mutants compared to WT at all stages of embryogenesis. Quantitative analysis of stage-wise cell volumetric data (mean with SEM) from a total of 1200 cells that were 3D rendered. N=150 cells from three embryos of each genotype at each stage. WT: Mean ± SEM— St. 11: 42.13 ± 0.93 µm^3^; St. 12: 53.81 ± 1.46 µm^3^; St. 13/14: 75.39 ± 1.73 µm^3^; and St. 15/16: 98.44 ± 1.86 µm^3^. *rib* null: Mean ± SEM—St. 11: 24.73 ± 0.65 µm^3^; St. 12: 48.43 ± 0.95 µm^3^; St. 13/14: 49.33 ± 1.29 µm^3^; and St. 15/16: 63.05 ± 1.63 µm^3^ **(E)** <u>Top:</u> WT and *rib* mutant SG cell volume during early stages (11 & 12) of tubulogenesis (P<0.05; two-tailed unpaired t-test; median and quartiles). <u>Bottom:</u> WT and *rib* mutant SG cell volume during late stages (13 – 16) of tubulogenesis (P<0.05; two-tailed unpaired t-test; median and quartiles). **(F)** α-<u>SAS</u> staining of the luminal membrane (arrowhead) reveals the tube elongation failure that is associated with the cell growth deficit in the *rib* mutant SG compared to the stage-matched control. Scale bar: 50 µm. **(G)** SG nuclear DNA volume is not affected by loss of *rib*. <u>Left:</u> 3D rendering of DAPI staining showed no difference in nuclear DNA volume between WT and *rib* mutant SGs. A total of 300 nuclei were 3D rendered, n = 150 from three embryos of each genotype from the late stages (15 – 16) of tubulogenesis. Scale bars: 10 µm. <u>Right:</u> Quantitative analysis of DNA volume in WT and *rib* mutants (two-tailed, unpaired t-test; median and quartiles). N.S.—Not significant. **(H)** Whole embryo volume is unchanged in *rib* mutants. <u>Left:</u> 3D rendering of whole embryos. Scale bar: 50 µm. <u>Right:</u> Quantitative analysis of embryonic volume showed no difference in mean volume of *rib* mutants versus WT. (n=5 embryos/group; two-tailed, unpaired t-test; mean with SEM). N.S.—Not significant.

SGs are specified by the Hox protein Sex combs reduced (Scr) (Andrew et al., 1994; Panzer et al., 1992), working with Extradenticle (Exd) and Homothorax (Hth) (Henderson and Andrew, 2000). Scr, Exd and Hth activate a core set of transcription factors—Fork head (Fkh), Salivary gland-expressed bHLH (Sage), Cyclic-AMP response element binding protein A (CrebA), Senseless (Sens), and Huckebein (Hkb)—the actions of which regulate diverse cell physiological processes during SG tubulogenesis, while maintaining cell fate and priming secretory function (Abrams and Andrew, 2005; Abrams et al., 2006; Fox et al., 2010; Fox et al., 2013; Johnson et al., 2020; Maruyama et al., 2011; Myat and Andrew, 2000a, 2002). A subset of these factors also controls the morphogenetic attributes of the SG tubulogenic program, *e.g.*, cell invagination (Fkh) and tube elongation (Hkb) (Chung et al., 2017; Myat and Andrew, 2000a, b, 2002).

The contributions of yet another nuclear factor—Ribbon (Rib)—to the SG morphogenetic program are less well understood (Bradley and Andrew, 2001; Xu et al., 2011). *rib* encodes a nuclear protein implicated in epithelial cell shape change in several tubular organs, including the embryonic SG, trachea, hindgut, and Malpighian tubules (Bradley and Andrew, 2001; Jack and Myette, 1997; Shim et al., 2001). Morphogenesis of non-tubular tissues, including the amnioserosa and somatic gonad precursors, is also affected by *rib* loss (Jack and Myette, 1997; Silva et al., 2016). The most striking loss-of-function *rib* phenotype in several organs is incomplete/failed tube elongation. Also, the cells lacking Rib function assume a more rounded or cuboidal shape in tissues where they normally form wedge and columnar shapes (Jack and Myette, 1997; Loganathan et al., 2016). In several of these tissues, including the SG, Rib has been shown to function tissue-autonomously (Bradley and Andrew, 2001; Silva et al., 2016).

Candidate-gene-based approaches have identified both cytoskeletal and membrane-localized proteins, including Crumbs (Crb), Rab11, and Moesin (Moe), as mediators of Rib-directed SG and tracheal tube elongation (Kerman et al., 2008). These results suggest that Rib directs tube elongation by facilitating apical membrane expansion/growth. Computational analysis of SG tube elongation, predicated on viscoelastic models, confirmed membrane expansion/growth as a critical feature compromised in *rib* mutants (Cheshire et al., 2008). The tube elongation defect in *rib* mutant SGs occurs without affecting specification, apicobasal polarity determination, junctional integrity, or cell number. A snapshot of cell volume at early stages in the *rib* mutant SG defined the tube elongation defect in terms of a cell growth/size defect, as it was significantly decreased compared to WT controls (Loganathan et al., 2016). Overall, these prior findings show that Rib controls multiple critical aspects of SG morphogenesis and suggest that cell growth may be a fundamental characteristic of Rib-mediated tube elongation.

Despite its confirmed role in tube elongation and association with apical membrane expansion/growth, the extent to which Rib affects the morphogenetic trajectory of SG tubulogenesis is unclear. Importantly, to our knowledge, cell growth as an integral component of organogenesis during the embryonic stages in *Drosophila* or other organisms, has not explicitly been considered. How does Rib affect SG cell growth? Is there a growth component interwoven into other morphogenetic processes, *e.g.*, cell invagination, apical membrane expansion, and collective cell migration, to facilitate SG tube elongation? To address these questions, we focused on potential transcriptional targets of Rib that were identified through an unbiased, genome-wide approach. We report that Rib binds to and upregulates ribosomal protein genes (RPGs) to drive cell growth (volume gain) as an integral component of embryonic SG morphogenesis. In addition to RPGs, Rib binds chaperones and translation factors required for protein synthesis suggesting augmented translation capacity as the basis for SG tubulogenic growth—an adaptation that likely facilitates SG function as a dedicated secretory organ. We find that Rib physically interacts with known regulators of RPG transcription to gain context-specificity for RPG promoter binding because its direct binding to DNA (in vitro) is of low affinity and specificity. Finally, we provide evidence that Rib binding to RPGs is specific for SG tube elongation, because the likely direct targets of Rib that mediate cell growth in yet another embryonic tube, the trachea, do not include the RPG repertoire. Collectively, these results identify early cell growth as a potential contributor to tube elongation and suggest that Rib activates cell growth programs that are customized for tissue function.

## RESULTS

### SG secretory cell growth is through Rib-dependent cytoplasmic volume gain

Loss of *rib* function results in a significant decrease in embryonic SG tube elongation without affecting secretory cell number suggesting that Rib is relevant to cell growth in tube elongation independently of cell division or death (Loganathan et al., 2016). To quantify individual cell size in both WT and *rib* mutant SG cells, we segmented images of secretory cells stained with α-MIPP1 (Multiple inositol polyphosphate phosphatase 1; a membrane marker; (Cheng and Andrew, 2015) from early and late stages of tubulogenesis (**Fig. 1C**). To allow for complete 3D cell surface rendering, only SG cells showing uniform membrane staining were included for volumetric analysis. The results revealed that WT SGs undergo a gradual increase in mean cell volume, discernible as a progressive, stage-wise growth series that more than doubled cell size over a period of about six hours (**Fig. 1D**). In contrast, developmental growth of *rib* mutant SG cells was significantly impaired—with both lower starting cell volumes (when SG cell markers are first discernable) and lower volume gains compared to WT (**Fig. 1D**). By late embryogenesis, *rib* mutant SG cells show a 36% reduction in mean overall size (**Fig. 1E**) and a shortened lumen as the tube fails to fully elongate (**Fig. 1F**).

The cell growth defect observed in *rib* mutant SGs is not due to impaired DNA amplification, *i.e.*, polytenization, because the mean DNA volume of *rib* mutant SG cells from St. 15/16—well beyond their first and only embryonic-stage endocycle at St.12 (Smith and Orr-Weaver, 1991)—was not different from WT (**Fig. 1G**). To ask if the growth loss in *rib* mutant SG cells is a bystander effect, *i.e.*, a co-occurrence or consequence of systemic growth deficiency at the level of the whole embryo, we performed whole embryo volumetric analysis at terminal stages of embryogenesis and found it was no different between WT and *rib* mutants (**Fig. 1H**).

Collectively, these results reveal that post-mitotic cell growth is integral to embryonic SG tubulogenesis. Rib-dependent SG secretory cell growth operates independently of whole embryo size determinants, and is concomitant with morphogenetic processes—*i.e.* invagination, shape change, and collective migration. SG cell growth manifests as a gradual Rib-dependent increase in cytoplasmic volume gain during embryonic-stage epithelial tubulogenesis.

### Rib binds ribosomal genes as well as morphogenetic effectors

*rib* encodes *a* 661-residue protein with two protein-protein interaction domains—an N-terminal BTB domain and a C-terminal coiled-coil domain—and a bipartite nuclear localization sequence that is partly embedded in a DNA binding domain homologous to that of the Pipsqueak (PSQ) transcription factor (**Fig. 2A**). Rib is dynamically expressed in all three germ layers with slightly higher levels observed in several epithelial tubular organs, *e.g.*, the SG and trachea, and in the mesoderm (**Fig. 2B**). Rib is nuclear, but is absent from the DAPI-concentrated regions of the nucleus that correspond to the nucleolus—the site of ribosome biogenesis (Pederson, 2011) (**Fig. 2C**).

**Figure 2.**
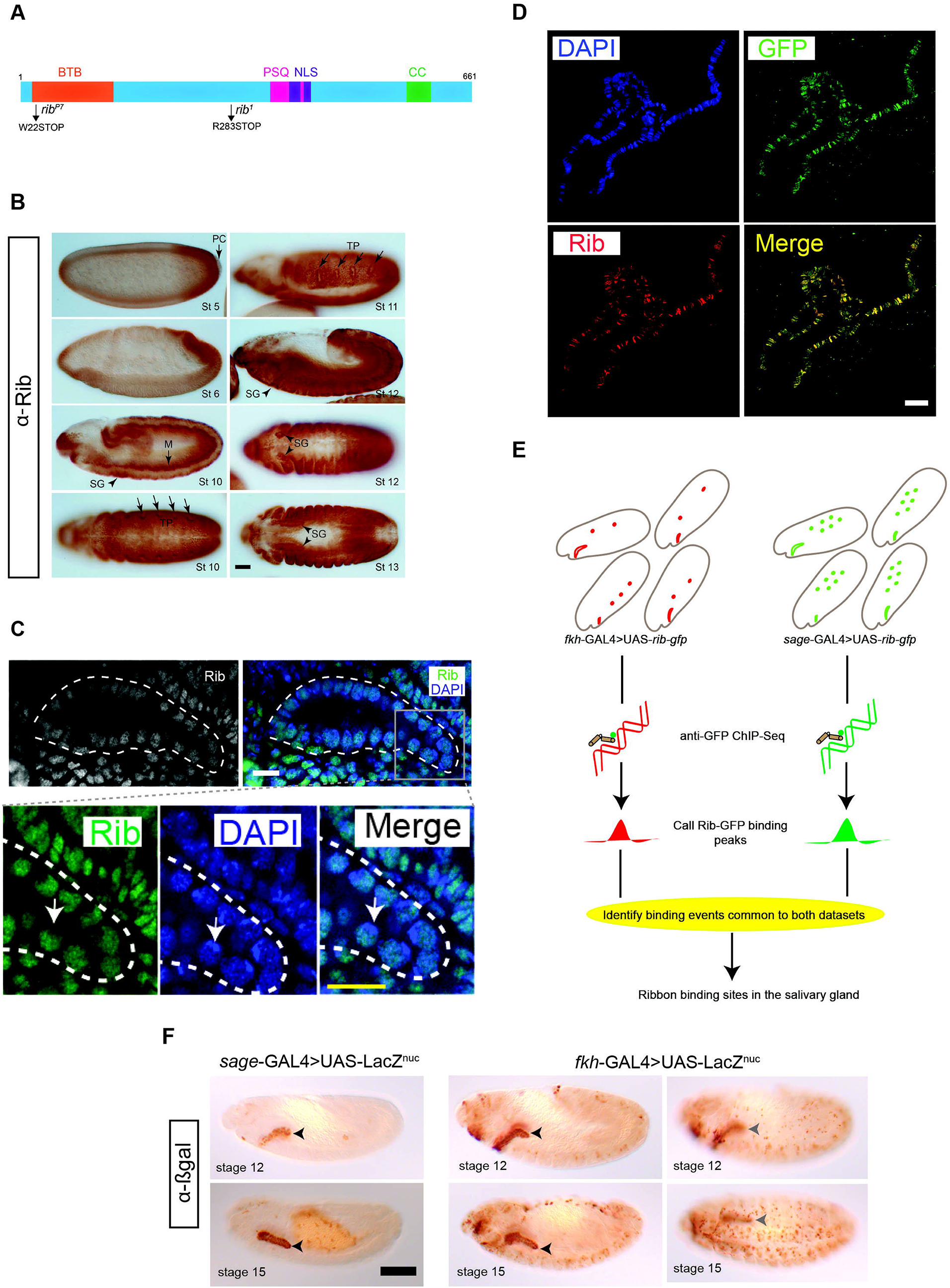
Strategy for identifying the transcriptional targets of Rib in the SG. **(A)** Rib protein diagram with the N-terminal Bric à Brac, Tramtrack, Broad (BTB) domain, the pipsqueak (PSQ) DNA-binding domain, the bipartite nuclear localization sequence (NLS) and a predicted coiled coil (CC) domain. Compound heterozygotes from the null alleles of *rib*—*rib^P7^* and *rib^1^*—were used throughout this study. **(B)** Rib staining in all three germ layers during mid-through late-embryogenesis. SG – salivary gland, M – mesoderm, PC – pole cells, TP – tracheal primordia. Scale bar: 50 µm. **(C)** Ribbon is nuclear. <u>Top:</u> White outline marks the SG. <u>Bottom:</u> Rib staining is absent in the DAPI-intense areas (nucleoli; arrow). Scale bar: 10 µm. **(D)** Rib-GFP, used for tissue-specific ChIP-Seq analysis, localizes to SG polytene chromosomes of L3 larvae, with a one-to-one correspondence of bands detected with anti-GFP and anti-Rib antisera. (Images from *sage*-GAL4>UAS-*rib-GFP*.) Scale bar: 10 µm. **(E)** Strategy for ChIP-Seq using *fkh*-GAL4 or *sage*-GAL4 to drive UAS-Rib-GFP expression. Tissue expression patterns of Rib-GFP in *fkh*-GAL4>UAS-*rib-GFP* embryos and *sage*-GAL4>UAS-*rib-GFP* embryos are illustrated in red (SG and hemocytes) and green (SG and midgut), respectively. SG is the only tissue common to Rib-GFP expression from both the drivers, therefore, an overlap of binding peaks from both driver datasets enriches for high-confidence, SG-specific Rib binding. **(F)** Expression domains of SG drivers—used in the Rib-GFP ChIP-Seq experiments—are indicated with nuclear βgal during early (stage 12) and late (stage 15) tubulogenesis. For the *fkh*-GAL4 driver, a different focal plane (right images) captures expression in mesodermally-derived cells. Arrowheads mark the SG. Scale bar: 50 µm.

To determine how Rib mediates SG cell growth, we re-examined previously generated SG-specific ChIP-Seq data obtained using a functional Rib-GFP that both rescued the tube elongation defect of *rib* mutants (Loganathan et al., 2016) and localized to distinct SG chromosomal loci recognized by both anti-GFP and anti-Rib antisera (**Fig. 2D**). To enrich for SG-specific Rib targets, we used binding data from experiments carried out with two different SG drivers—*sage*-GAL4 and *fkh*-GAL4—whose expression domains overlap in only the SG (**Fig. 2E, F**). Each experiment was done with two biological samples utilizing a GFP antibody that has been robustly tested for ChIP-Seq applications (Kudron et al., 2018; Sin et al., 2017). We used a publicly available ChIP-Seq pipeline from ENCODE that utilizes Irreproducible Discovery Rate (IDR) correction on replicates (Li et al., 2011) to identify robust binding peaks from each experiment and filtered to retain only those peaks found with both SG drivers; this yielded a high-confidence, highly correlated set of 436 SG-specific Rib binding peaks (**Fig. 3A**).

**Figure 3.**
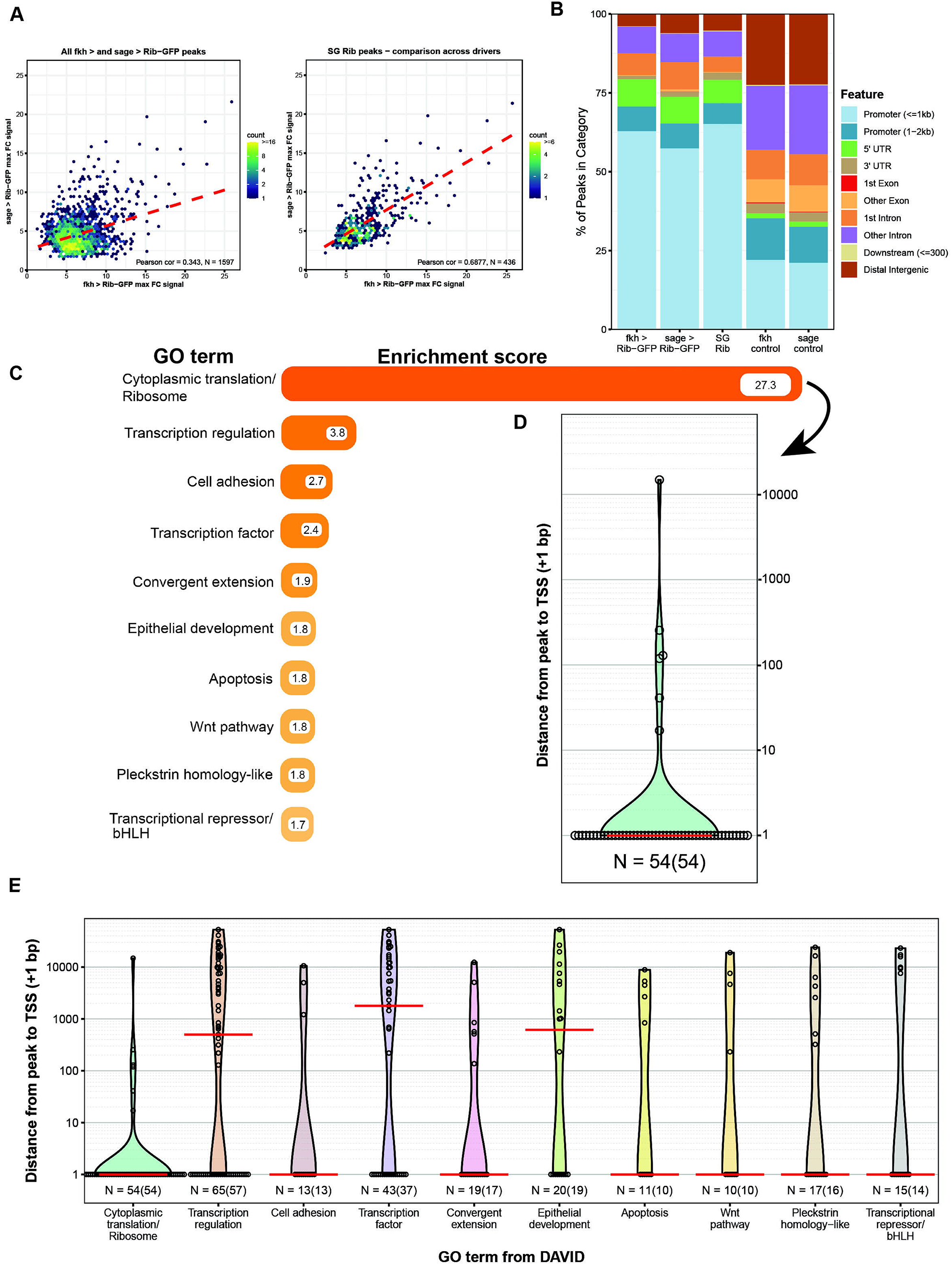
Analysis of salivary gland Rib-binding data implicates ribosome/translation. **(A)** <u>Left plot:</u> Rib-binding signal enrichment comparison between the GAL4 drivers, *i.e.*, all IDR peaks from *fkh*-GAL4>UAS-*rib-GFP* and *sage*-GAL4>UAS-*rib-GFP*. <u>Right plot:</u> Rib-binding signal enrichment comparison across the tissue-specific GAL4 drivers, *i.e.*, overlapping regions of the IDR peaks from the *fkh*-GAL4>UAS-*rib-GFP* and *sage*-GAL4>UAS-*rib-GFP* experiments, showing a high positive correlation—high confidence Rib-binding for SG-specific targets. **(B)** Binding signal feature analysis in both driver experiments show that the Rib binding peaks (≈ 75%) localize in the vicinity of target gene promoters, especially in the SG Rib peaks. Controls indicate expected distribution of binding events if they were random with respect to gene elements. **(C)** Functional clustering of Rib-bound SG genes under gene ontology categories (GO terms) according to the Database for Annotation, Visualization and Integrated Discovery (DAVID). The top ten GO terms with their associated enrichment scores are shown. The predominant category “Ribosome” represents the binding of Rib to RPGs with a nearly seven-fold greater enrichment compared to the next-ranked category. *See* ***Table S1*** *for meta data*. **(D)** Rib binds to the transcription start sites (TSS) of RPGs. Open circles indicate the distance between the Rib binding peak and the TSS for individual genes (+1 bp). Red line indicates the median distance from the peak to TSS (+1 bp). **(E)** Enrichment of TSS-proximal Rib binding across the top ten functional classes. Including the ribosome, seven of the top ten enriched GO classes primarily show Rib binding to the TSS. Open circles indicate the distance between the Rib binding peak and the TSS for individual genes (+1 bp). Red lines indicate the median distance from the peak to TSS (+1 bp). N = number of peaks versus (number of genes).

Assessment of the global features of Rib binding in the SG showed a high propensity, *i.e.*, > 60%, for promoter-proximal binding of Rib with both experiments (**Fig. 3B**). Application of the Database for Annotation, Visualization and Integrated Discovery (DAVID) gene ontology program for the 413 genes that correspond to the 436 overlapping SG Rib binding peaks revealed the ribosome as the primary Rib target (**Fig. 3C – D, Table S1**), with a nearly seven-fold higher enrichment score compared to the second-ranked target gene cluster. Other top clusters include transcription, cell adhesion and convergent extension. Taken together, these results are consistent with the purported function of Rib as a mediator of tissue morphogenesis (Bradley and Andrew, 2001; Jack and Myette, 1997; Loganathan et al., 2016). They also suggest a plausible mechanism for Rib’s role as a regulator of SG cell growth targeting the ribosome and, consequently, cellular translational efficacy, as motivated by similar findings of a causal link between ribosomal transcriptional regulation and cell size in a relatively simple model system, *S. Cerevisiae* (Jorgensen et al., 2004). Interestingly, the majority of the top ten SG target clusters (7/10) featured promoter-proximal Rib binding, with Rib binding at the transcription start site (TSS) for the majority of genes within each of seven clusters (**Fig. 3E**).

### Rib binds the TSS of RPGs for their regulation

The ribosome is composed of two genomically-encoded components—ribosomal RNAs (rRNAs) and ribosomal proteins. Examination of the gene cluster encapsulated by the gene ontology term *ribosome/cytoplasmic translation* revealed that Rib binds RPGs and not the genes encoding rRNAs. Visualization of Rib binding at SG-expressed RPGs (**Table S2**) using the Integrative Genomics Viewer (IGV) revealed that 73 of the 84 SG-expressed RPGs are bound by Rib, *i.e.*, 87% coverage, with peaks exceeding a Log_10_ binding likelihood threshold ≥ 4. Among the 73 Rib-bound RPGs, 64 show Rib-bound peaks at or above this threshold from both driver datasets, seven have such peaks in the ‘*fkh*-GAL4 ChIP-Seq only’ dataset and two have such peaks in the ‘*sage*-GAL4 ChIP-Seq only’ dataset (**Fig. 4A**). In every case, the Rib-bound region on chromatin spans the RPG TSS (**Fig. 4B – C, S1A – S1E**). Even in the class of RPGs classified as ‘Rib-not bound,’ the strongest relative Rib-binding signal almost always spans the TSS of the corresponding RPG (**Fig. 4D, S1A – S1E**). Rib-bound RPG loci are not confined to a specific chromosome (**Fig. S2A**). Neither do the Rib-bound RPGs show a preferential bias for a particular ribosomal subunit (large or small) or any specific domain within a subunit (**Fig. S2B**). Thus, we surmise that Rib likely binds all SG-expressed RPGs, but that the technical limitations of the ChIP-Seq approach prevented absolute recovery of all RPG binding events.

**Figure 4.**
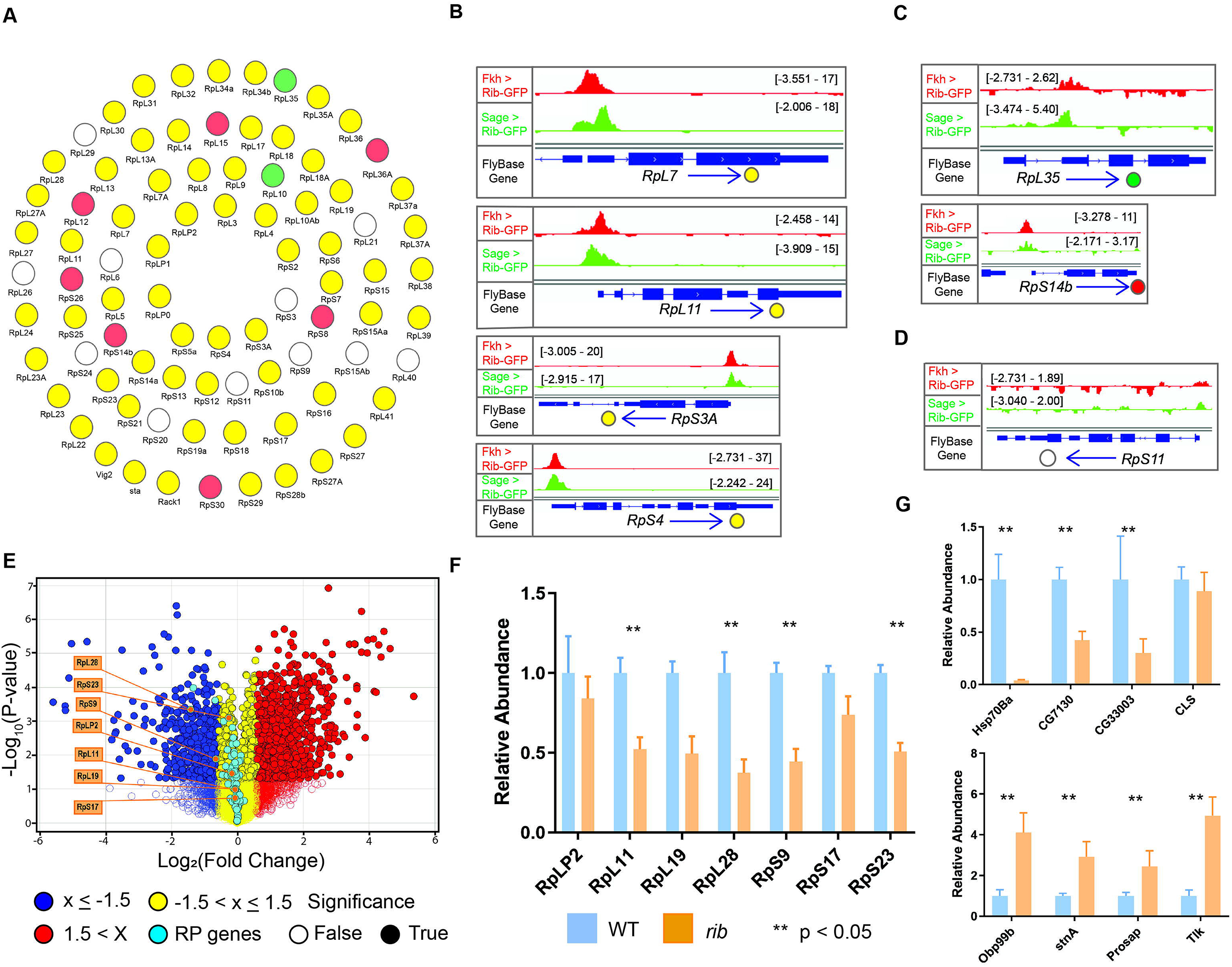
Rib immunoprecipitates RPGs and is required for their full levels of expression. **(A)** Schematic summary of Rib binding to SG-expressed RPG promoters. Filled circles represent genes bound by Rib (Log_10_ binding likelihood threshold ≥ 4). Unfilled circles represent genes not bound by Rib (Log_10_ binding likelihood threshold < 4). Yellow: set of genes bound by Rib and above threshold in both *fkh*-GAL4 and *sage*-GAL4 experiments; Red: set of genes bound by Rib and above threshold in the *fkh*-GAL4 experiment only; Green: set of genes bound by Rib and above threshold in the *sage*-GAL4 experiment only. **(B)** Representative binding profiles of Rib-GFP for genes with peaks from both datasets (*fkh*-GAL4>UAS-*rib-GFP* and *sage*-GAL4>UAS-*rib-GFP*): *RpL7*, *RpL11*, *RpS3A*, and *RpS4*. Binding profiles from the *fkh*-GAL4 and the *sage*-GAL4 datasets are shown by red and green tracks, respectively. Signal intensity range for the regions shown are in brackets*. See* ***Fig. S1 A – E*** *for all RpG binding profiles*. **(C)** Representative binding profiles of Rib-GFP for genes with peaks (Log_10_ binding likelihood threshold ≥ 4) with only one of the datasets: *RpL35* (*sage*-GAL4>UAS-*rib-GFP*) and *RpS14b* (*fkh*-GAL4>UAS-*rib-GFP*). Note that although the alternate track did not meet the binding threshold (≥ 4), its signal peak overlaps with the signal peak that did reach threshold. **(D)** Representative signal profile of Rib-GFP on a RPG from Rib-not-bound category (Log_10_ binding likelihood < 4 in both datasets): *RpS11*. Note that although neither track met the binding threshold (≥ 4), the highest signal, nonetheless, spans the TSS. **(E)** Volcano plot of whole-embryo microarray gene expression analysis shows genes that were downregulated (blue) or upregulated (red) at least 1.5-fold in *rib* null mutants compared to WT. Transcripts with fold-change values between −1.5 and 1.5 are shown in yellow. Unfilled circles represent transcripts with fold change values that were not statistically significant between the groups, *i.e.*, P > 0.05. RPGs are highlighted by cyan circles. A majority of RPG transcripts were downregulated in the *rib* mutants compared with WT. Named RPGs (orange) belong to the subset whose levels were also examined by RT-qPCR analysis. *See* ***Table S2*** *for metadata*. **(F)** RT-qPCR results from whole embryo transcripts for several RPGs show reduced expression levels in *rib* mutants compared to the WT, with most showing a significant decrease. **P<0.05; Mann-Whitney *U* test. **(G)** Non-ribosomal Rib targets show both decreased (top) and increased (bottom) expression in *rib* mutants compared to the WT. (*Data from Loganathan et al., 2016)* **P<0.05; Mann-Whitney *U* test.

To determine if RPGs are transcriptionally regulated by Rib, we first reexamined the whole-embryo microarray analysis comparing gene expression levels in WT to *rib* mutants in mid-to late-stage embryogenesis (Loganathan et al., 2016). A majority of RPGs, *i.e.*, 67/84, are downregulated in *rib* mutants compared to WT (**Fig. 4E, Table S2**). Although merely 4/67 RPGs (two Rib-bound and two Rib-not-bound) show a greater-than 1.5-fold downregulation in *rib* mutants, the overall trend—notwithstanding the specific fold-change values—suggests a dependence of RPGs on Rib for their full transcriptional activation. Indeed, both Rib-bound and Rib-not-bound RPGs depend on Rib for transcriptional upregulation, with 59/67 RPGs whose expression went down in *rib* mutants belonging to the Rib-bound category and eight belonging to the Rib-not-bound category (**Table S2**).

We next tested a subset of RPGs from both the large and small ribosomal subunits for Rib-dependent transcriptional regulation using RT-qPCR analysis of whole embryos. The subset included six Rib-bound genes— RpLP2, RpL11, RpL19, RpL28, RpS17, and RpS23—and one Rib-not-bound gene—RpS9. The levels of all seven transcripts were lower in *rib* mutants compared to WT, with four of the tested RPGs showing significantly reduced levels of expression (**Fig. 4F, Table S3**). In contrast, RT-qPCR analysis of non-ribosomal genes bound by Rib (using the same RNA samples) revealed both increased and decreased levels of expression in *rib* mutants compared to WT (**Fig. 4G**). The RT-qPCR results, hence, support a requirement for Rib to boost RPG transcription to WT levels in the embryo since, although transcript levels are decreased in *rib* mutants compared to WT, RPG expression is, nonetheless, still observed.

### Rib transcriptionally upregulates SG RPGs and *rib* loss adversely impacts markers of ribosome biogenesis and cell translation

The abundance of RPG transcripts in the SG relative to other tissues during embryogenesis (http://fly-fish.ccbr.utoronto.ca/) prompted us to assay the effects of *rib* loss on RPG transcript levels using fluorescent *in situ* hybridization (FISH). Transcript levels for all five of the RPGs we tested were notably reduced in *rib* mutants compared to WT, supporting the results from our RT-qPCR experiments (**Fig. 5A, S3**). Interestingly, the FISH experiments revealed not only a decrease in RPG levels in the SG but also in the mesodermally-derived cells surrounding the SG, an unexpected finding that is nonetheless consistent with the high levels of Rib also detected in the embryonic mesoderm and its derivatives (**Fig. 2B – C**). Restoration of Rib function using *fkh*-GAL4, which drives UAS-transgene expression in the SG and in a subset of mesodermally-derived cells (**Fig. 2F**), was sufficient to rescue RPG transcript levels in *rib* mutants. The residual level of RPG transcript (and presumably protein) in *rib* mutant SGs may suffice to meet the basal translation requirements since *rib* mutant SG cells do not die (Loganathan et al., 2016), but may fall short of fulfilling the increased translational capacity demanded by morphogenetic growth and/or the increased secretory output of this organ.

**Figure 5.**
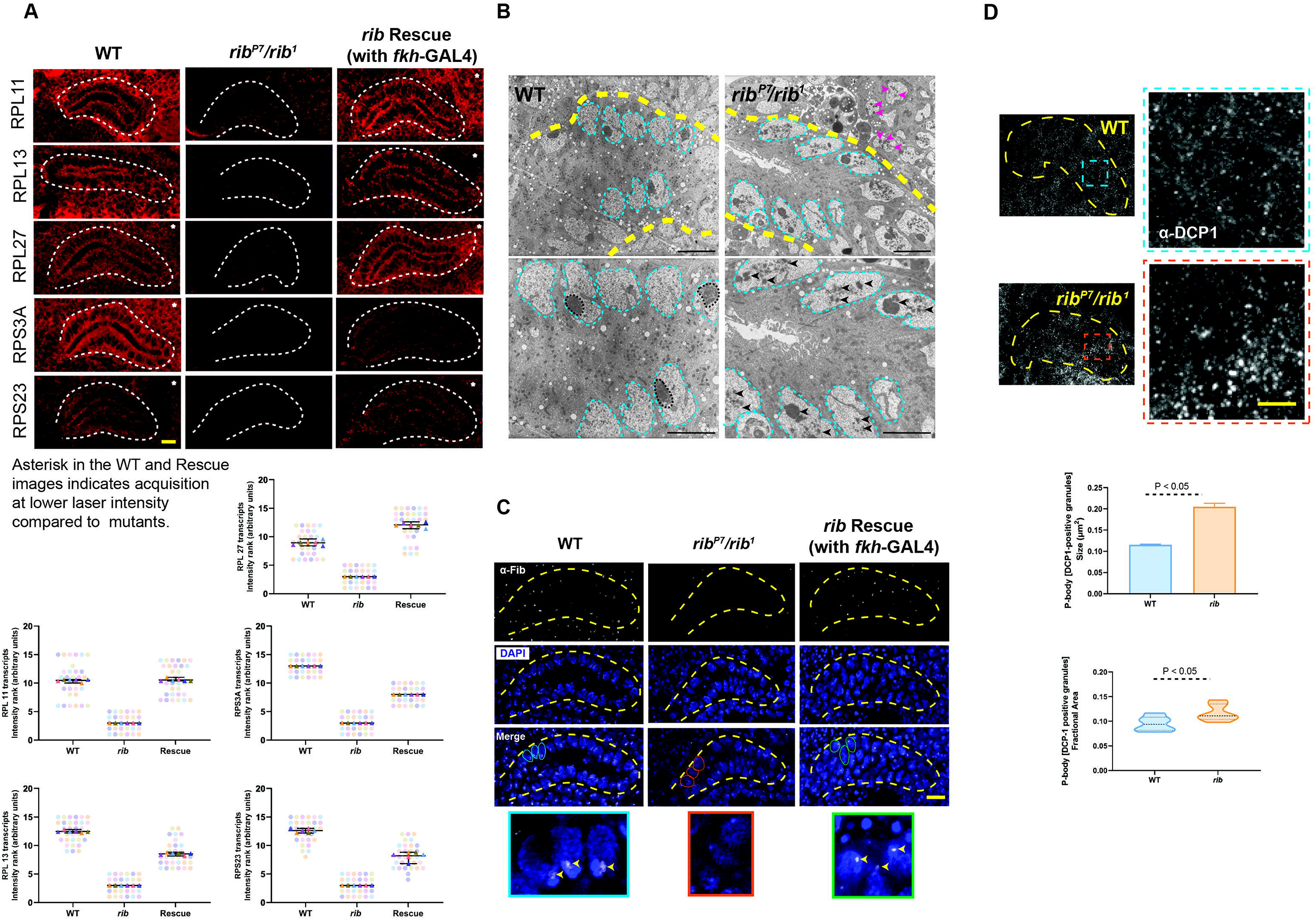
*rib* is required for full levels of RPG expression and *rib* mutants show defects consistent with ribosome deficiency. **(A)** <u>Top:</u> Representative images from fluorescent *in situ* hybridization (FISH) of RPGs reveal high level expression in the WT SG (white outline) and a significant reduction in *rib* mutants, which is rescued by *fkh*-GAL4 driven expression of *rib*. <u>Bottom:</u> SuperPlots showing the scores assigned by six observers ranking RPG transcript signal intensities in 15 blinded samples in the FISH experiments. Note that the observers consistently ranked what turned out to be the *rib* null samples as having the lowest SG signal intensity. Scale bar: 10 µm. *See* ***Fig. S3*** *for DAPI channel*. **(B)** Transmission electron micrographs from sections of embryonic SGs (yellow outline) revealed aberrant nucleolar morphologies in the *rib* mutants compared to WT. (Images in the bottom row are slightly higher magnifications.) Whereas the nuclear morphologies (cyan outline) are comparable between the groups, the electron-dense compaction of nucleolar condensates in the WT (black outline) is rarely observed in the *rib* mutants with the majority of mutant cells showing fragmented or dispersed morphology characteristic of the nucleolar decondensation phenotype (black arrowheads). Adjoining mesodermal cell nucleoli also have a fragmented or dispersed morphology (magenta arrowheads) in the *rib* mutant. Scale bar: 2 µm. **(C)** Fibrillarin-positive nucleolar punctae observed in WT are lost in *rib* mutants (yellow outline—SG); *fkh-*GAL4 driven expression of *UAS-rib* was sufficient to rescue its localization in SGs and in a subset of surrounding mesoderm-derived cells that express *fkh-*GAL4. A subset of outlined nuclei (WT—cyan; *rib* mutant—orange; and rescue—green) in the “Merge” row is enlarged in the bottom row to highlight the nucleolar Fibrillarin (arrowheads). Scale bar: 10 µm. **(D)** P body granules (α-DCP1 staining), indicators of untranslated mRNA accumulation, are larger and relatively more abundant in *rib* mutants than in WT. <u>Top</u>: SG (yellow outline) and the enlarged region of interest (blue/orange boxes) are shown. Scale bar: 2 µm. <u>Bottom:</u> Quantitative analysis showed a significant increase in the P body granule size in *rib* mutant SG cells (0.205 ± 0.008 µm^2^) compared to WT (0.115 ± 0.001 µm^2^) (Unpaired t-test with two-tailed P value <0.001; mean ± SEM; n = 6 SGs/group). Quantitative analysis of P body granule fractional area also showed a significant increase in their relative abundance in *rib* mutant SG cells (0.117 ± 0.007) compared to WT (0.0951 ± 0.006) (Unpaired t-test with two-tailed P value: 0.05; median and quartiles; n = 6 SGs/group).

To learn if the decrease in RPG transcript levels in *rib* mutant cells is associated with changes in markers of ribosome biogenesis and translation, we examined SG nucleoli and RNA processing bodies (P bodies). Nucleoli are the sites of rRNA processing and ribosome subunit assembly, for which stoichiometric levels of RPs are an absolute prerequisite (Pederson, 2011). Decreased availability of RPs could perturb nucleolar homeostasis and impacts its morphology. Accordingly, our TEM analysis showed the WT SG nucleolus as a single compact electron-dense condensate within each nucleus. In the majority of *rib* mutant cells, however, the nucleolus was both highly fragmented and widely dispersed (**Fig. 5B**). This aberrant morphology is reminiscent of the nucleolar stress/decondensation phenotypes typically associated with abnormal ribosome biogenesis and decreased cell growth (Marinho et al., 2011). We also observed loss of staining for Fibrillarin—a rRNA 2’-O-methyltransferase required for pre-rRNA processing—in *rib* mutant SG nucleoli (**Fig. 5C**). Fibrillarin localization within nucleoli is indicative of ribosome biogenesis and growth in multiple cell types (Baker, 2013; Sriskanthadevan-Pirahas et al., 2018a; Sriskanthadevan-Pirahas et al., 2018b). Importantly, expression of UAS-*rib* using the *fkh*-GAL4 driver was sufficient to rescue the Fibrillarin nucleolar staining, further supporting a role for Rib in ribosome biogenesis. We also observed changes in the nucleolar morphology and Fibrillarin staining of some mesodermally-derived cells (**Fig. 5B – C**), concordant with the decrease in their RPG transcript levels on FISH. The effects of Rib on nucleolar morphology are likely an indirect consequence of reduced RP levels since Rib itself does not localize to nucleoli (**Fig. 2C**) and neither does it bind *Fibrillarin* (SG ChIP-seq). We cannot, however, exclude the possibility that Rib binds and regulates some other critical but unknown nucleolar component.

We investigated the impact of *rib* loss on translation by comparing P body size in *rib* mutant and WT SG cells. P bodies are aggregates of untranslated mRNPs associated with translational repression and their size is proportional to the amount of untranslated mRNAs (Parker and Sheth, 2007; Teixeira et al., 2005). Staining for Decapping Protein 1 (DCP1), a core component of the P body-associated decapping machinery (Ingelfinger et al., 2002), revealed a significant, nearly two-fold increase in the aggregate size of P bodies in *rib* mutants (**Fig. 5D**). The fractional area of SG P body granules (total aggregate area/gland area) was also significantly increased in the *rib* mutants compared to WT (**Fig. 5D**), suggesting a larger pool of untranslated mRNAs associated with the loss of *rib*. These results, in addition to confirming Rib’s proposed role in augmenting SG cell translational capacity, also indicate that the RPG transcript level decrease in *rib* mutants is not because of any generic decrease in levels of nuclear transcription. Taken together, these results support a role for Rib in boosting SG RPG transcription—a prerequisite for increased ribosome biogenesis and efficient translation of mRNAs—to prime the organ for its secretory function.

### Rib binding to DNA *in vitro* is direct, weak, and not sequence-specific

We next explored the basis of Rib binding to RPG DNA by using Multiple Em for Motif Elicitation (MEME) analysis (Bailey and Elkan, 1994) to identify potentially conserved sequence motifs. Using the RPG sequences immunoprecipitated by Rib, we identified several motifs, the top five of which are shown (**Fig. 6A**). Mapping the sequence motifs in the 200bp region centered around the TSS (+1) for the 73 Rib-bound and 11 Rib-not-bound RPGs revealed a striking pattern in their organization (**Fig. 6B**, **S4**). The top-ranked TC-rich sequence (**Fig. 6A – B**; red motif) spans the TSS in almost all of the RPGs—both Rib-bound and Rib-not-bound. The other motifs are present in only a subset of RPGs: Ohler Motif-1 (cyan) in 36/84 enhancers (Ohler et al., 2002); the motif with similarity to the vertebrate ETS-1 binding site (green) in 15/84 enhancers (Sharrocks, 2001; Sharrocks et al., 1997); the Dref binding motif (purple) in 32/84 enhancers (Hirose et al., 1993; Hirose et al., 1996); and the motif resembling an NFI/CTF-halfsite (yellow) in 8/84 enhancers (Elateri et al., 2003; Gronostajski, 2000). Moreover, the relative position of the motifs is also conserved across RPG promoters (**Fig. 6B**), with the Ohler Motif-1(Cyan) and NF1/CTF-halfsite (yellow) being the closest motifs upstream of the TSS/TC-rich motif (red), and the Dref consensus sites (purple) mapping further upstream. The ETS-1 like motif (green) always occurs downstream of the TSS. The lack of a distinguishing motif-feature between RPGs bound by Rib and those not bound by Rib supports our supposition that the lack of Rib binding for the 11 RPGs in the Rib-not-bound group (**Fig. 6B**, bottom) is likely a technical artifact from ChIP-Seq.

**Figure 6.**
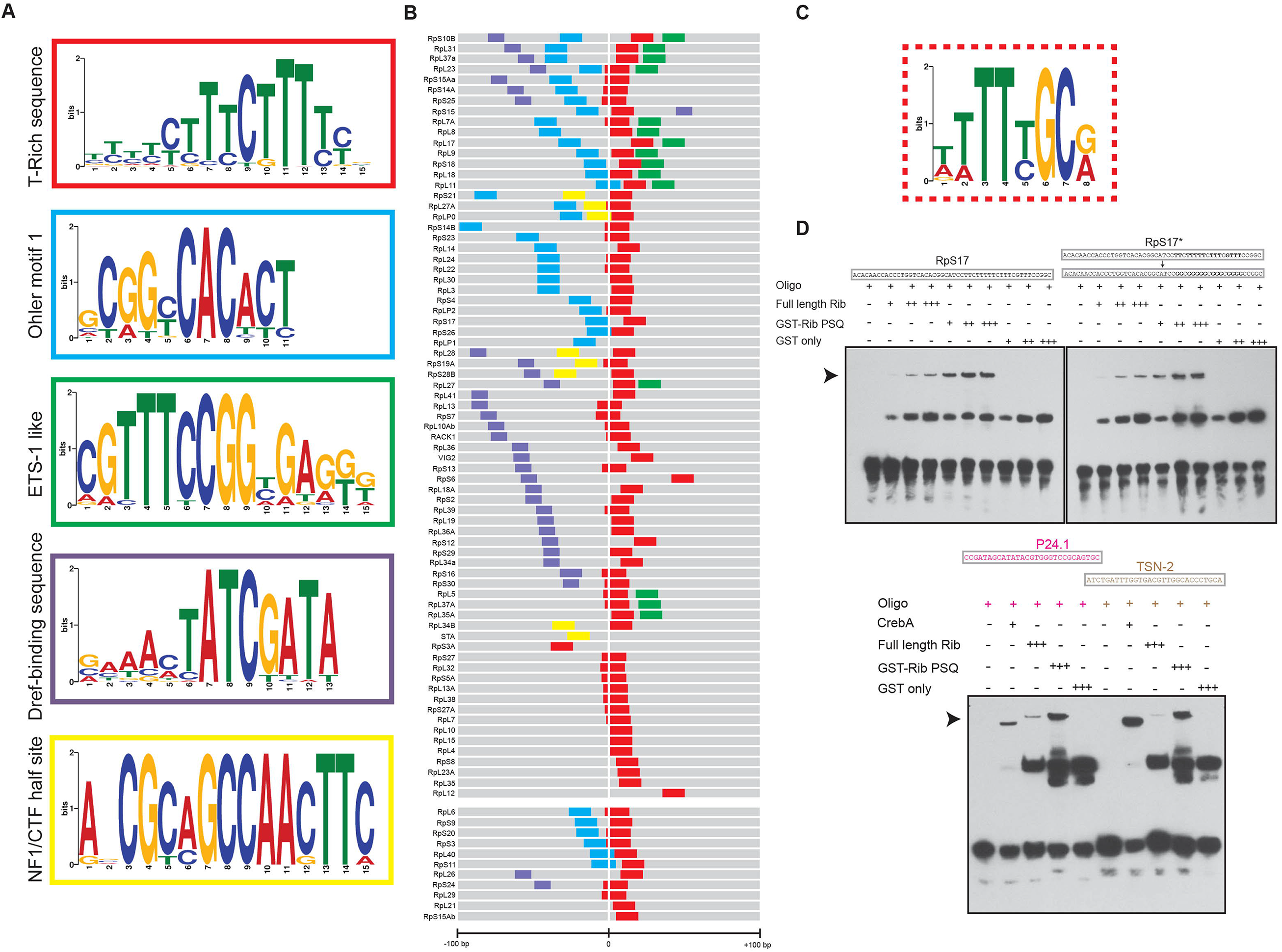
Rib pulls down sequences containing known RPG regulator motifs and binds to RPG enhancer DNA *in vitro* with low affinity and specificity. **(A)** Top five motifs identified by MEME-ChIP analysis of RPG sequences pulled down with Rib in the SG ChIP-Seq. **(B)** Representation of the enriched motifs from (**A**) embedded in the 200 nucleotides flanking the TSS of RPGs. The color code for motifs within putative enhancers is according to those used in the outlines of (**A**). TSS (+1) were obtained from FlyBase. The top block of enhancers (RpS10B through RpL12) includes genes bound by Rib and the bottom block (RpL6 through RpS15Ab) includes RPGs not bound by Rib in the ChIP-Seq analysis. *See* ***Fig. S4*** *for sequence lineup*. **(C)** T-rich motif ascertained for Rib binding from a bacterial one-hybrid analysis with the Rib PSQ domain (source: FlyFactorSurvey; URL: mccb.umassmed.edu/ffs). **(D)** EMSAs reveal that full-length Rib and its DNA-binding PSQ domain directly bind the RpS17 promoter in a concentration-dependent manner. <u>Left:</u> Rib-dependent mobility shift is indicated by the arrowhead. <u>Right:</u> EMSAs in which the T’s in the RpS17 promoter were replaced with G’s in the TC-rich motif of the RpS17* enhancer. <u>Bottom:</u> CrebA and Rib bind DNA containing the CrebA consensus sites in P24.1 and TSN-2—two bona fide transcriptional targets of CrebA. *See* ***Fig. S5*** *for EMSAs with several additional RPG TSS sequences.* All EMSAs were performed twice with identical results.

RPGs, as part of the housekeeping gene repertoire, feature promoter-proximal enhancers in contrast to the distal enhancers often associated with developmental genes (Zabidi et al., 2015). Three out of the top five consensus sequence motifs—TC-rich, Ohler Motif-1, and Dref motifs—that emerged from the DNA immunoprecipitated by Rib have, hitherto, been implicated in RPG transcription. The TC-rich sequence, abutting the TSS, harbors the TCT-core promoter region from which TATA box binding protein-related factor 2 (Trf2) initiates RPG transcription (Wang et al., 2014). Trf2, however, lacks sequence-specific DNA-binding activity, and therefore, likely relies on other transcription factors for its recruitment to the core promoter region. Two sequence-specific DNA-binding transcription factors— Motif 1 binding protein (M1BP) and DNA replication-related element factor (Dref)—bind to a subset of RPGs featuring their respective binding sites and likely recruit Trf2 to the core promoter region for initiating RPG transcription (Baumann and Gilmour, 2017; Hochheimer et al., 2002; Yamashita et al., 2007).

A previous study, utilizing a bacterial one-hybrid system-based screen, moreover, uncovered a preferred binding site for the DNA-binding PSQ domain of Rib as a T-rich sequence, not unlike the TC-rich consensus sequence highlighted by our MEME analysis of DNA immunoprecipitated by Rib, and that of the RPG core promoter region (**Fig. 6C**) (Noyes et al., 2008; Zhu et al., 2011). We, therefore, asked if purified full-length Rib or its PSQ DNA binding domain can directly bind RPG promoter-proximal enhancers *in vitro*. EMSAs with ten different RPG enhancers—all harboring the TC-rich promoter sequence and belonging to either categories, *i.e.*, Rib-bound or Rib-not-bound—revealed that both untagged full-length Rib and the GST-tagged DNA binding PSQ domain of Rib bind RPG enhancers in a concentration-dependent manner (**Fig. 6D, S5, Table S4**). The direct binding of full-length Rib or its PSQ domain to DNA is, however, not sequence-specific as revealed by (i) their ability to bind an altered RPG enhancer sequence in which the run of T’s was substituted by a run of G’s in the TC-rich promoter region (**Fig 6D**); and (ii) by the ability of Rib to shift DNA containing consensus binding motifs for CrebA from two genes, *P24-1* and *Tudor-SN*, that are not SG Rib-bound targets from ChIP-Seq (Fox et al., 2010; Johnson et al., 2020). In addition, Rib binds DNA with relatively weak affinity since the binding to the RPG enhancers required nearly 9x higher Rib protein concentration for electrophoretic mobility shifts as required for CrebA-dependent shifts of CrebA target sequences.

Also, the Rib PSQ domain bound DNA better than full-length Rib, suggesting some potential auto-inhibition with full-length Rib to DNA binding in vitro. In the ChIP-Seq experiment, Rib co-IPs with far fewer genes than observed for other *Drosophila* transcription factors (Acharya et al., 2012; Busser et al., 2012; Chambers et al., 2017). For example, CrebA exhibits high affinity sequence-specific binding *in vitro* and co-IPs with an order of magnitude more SG genes than Rib under nearly identical ChIP-Seq experimental conditions (Johnson et al., 2020).

Together, these results reveal that, on its own, Rib can bind DNA, but with poor affinity and virtually no sequence-specificity. Thus, other factors must contribute to the strength and specificity of Rib binding in the SG. Indeed, Rib-specific binding of RPGs to boost their expression could possibly be mediated, at least in part, through interactions of Rib with the sequence-specific binding proteins—MIBP and Dref—previously implicated in RPG regulation (Baumann and Gilmour, 2017; Hochheimer et al., 2002; Yamashita et al., 2007), and whose consensus motifs are harbored by RPG sequences that precipitated with Rib in the ChIP-Seq experiments. We also considered the possibility of Rib gaining context-specificity by interacting with Trf2, which initiates RPG transcription without directly binding the core TCT promoter (Wang et al., 2014).

### Rib interacts with known regulators of RPG transcription

Bolstering the possibility that known regulators of RPG transcription could both provide context and facilitate Rib binding to the RPG promoters, the developmental transcript expression profiles of Trf2, M1BP, and Dref trace trajectories closely in tandem with Rib, with all factors reaching their peak expression 4h – 10h post-fertilization, and dropping to near baseline levels at the end of embryogenesis (**Fig. 7A**). The timely increase in the expression of these RPG transcription factors, moreover, aptly precedes the peak levels of expression of RPG transcripts (**Fig. 7A**, inset), *i.e.*, 10h – 12h of embryogenesis.

**Figure 7.**
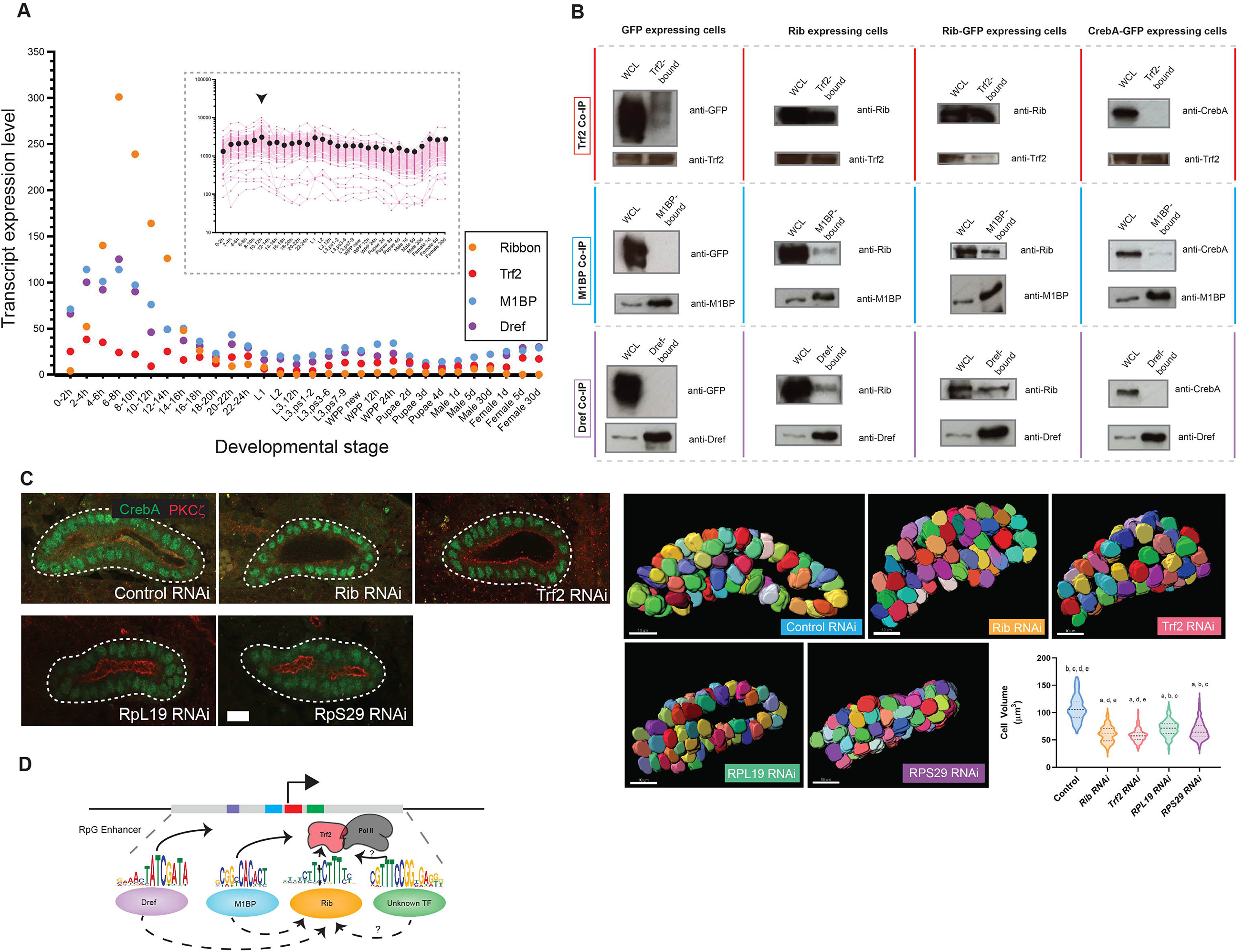
Rib co-immunoprecipitates with known transcriptional regulators of RPGs, and isolated RNAi knockdown of Rib-interacting RPG regulators or individual RPGs result in embryonic SG cell growth defects. **(A)** Transcript expression profiles of *rib*, *Trf2*, *M1BP*, and *Dref* during *Drosophila* development. The highest-level expression of all four genes occurs during embryonic stages corresponding with SG morphogenesis and cell growth. <u>Inset:</u> Transcript expression profiles for all 84 RPGs across developmental stages reveals the relatively high levels achieved during 10 – 12h of embryogenesis (arrowhead). Pink tracks—Individual RPGs; Black Dots—Median Values. Source data for the plots obtained from FlyBase (modENCODE temporal expression data). **(B)** Co-IP experiments were performed in S2R+ cells transfected with constructs expressing UAS-*gfp* (negative control), UAS-*rib*, UAS-*rib*-*gfp*, and UAS-*CrebA*-GFP (negative control), respectively. All three Co-IP fractions—Trf2-bound, M1BP-bound, and DREF-bound—contain both Rib and Rib-GFP. WCL: Whole Cell Lysate. Size (kDa)—Rib (≈ 70); CrebA (≈ 57); Trf2 short-form (≈ 75); M1BP (≈ 55); Dref (≈ 86); and GFP (≈ 30). A weak GFP signal was observed in the TRF2-bound fraction of GFP-expressing cells; and the weak CrebA signal observed in the M1BP-bound fraction of CrebA-GFP expressing cells was absent in its biological replicate experiment. **(C)** <u>Left:</u> Cell growth deficit is observed in SG RNAi knockdown of Rib, Trf2, and either of the two representative RPGs—RpL19 and RpS29—in St. 15, 16 embryos. <u>Right:</u> Quantitative analysis of cell volumetric data. A total of 750 cells from 15 SGs were 3D rendered. (n=150 cells/genotype; median and quartiles; Kruskal-Wallis test; significant difference indicators—P ≤ 0.05 from each group mean compared to (a) GFP RNAi control, (b) Rib RNAi, (c) Trf2 RNAi, (d) RpL19 RNAi, (e) RpS29 RNAi. Scale bar: 10 µm. **(D)** Model proposed for RPG regulation by Rib in the embryonic SG with a generic promoter-proximal RPG enhancer featuring putative binding sites for the multiple sequence-specific transcription factors that bind Rib and Trf2 to boost RPG transcription. Dashed arrows indicate the interaction of Rib (this study) with TRF2, M1BP, and Dref. Solid arrows indicate the interaction of Dref and M1BP with TRF2 (Hochheimer et al., 2002; Wang et al., 2014). Unknown or uncharacterized interactions are indicated by ?.

To test the possibility of physical interactions between Rib and the known regulators of RPG transcription in a relatively simplified system, we performed Co-IP experiments in S2R+ cells in which we drove expression of Rib, Rib-GFP, GFP only, or CrebA-GFP—the latter two serving as negative controls. S2R+ cells express endogenous TRF2, M1BP and Dref (FlyBase expression data). In the Co-IP experiments, we found that all three cell lysate fractions—TRF2-bound or M1BP-bound or Dref-bound—immunoprecipitated Rib as well as Rib-GFP, thus revealing the ability of Rib to physically interact with each of the three known activators of RPG transcription (**Fig. 7B**).

We next asked if knockdown of a likely Rib co-factor or of an RPG could phenocopy the cell growth deficit observed in *rib* mutant SGs. We quantified cell volume of St. 15,16 SG secretory cells in embryos expressing RNAi constructs targeting Rib or Trf2 or RpL19 or RpS29 under the control of the *fkh*-GAL4 driver. Knockdowns of Rib or Trf2 or RpL19 or RpS29 showed significant cell size deficits compared to the GFP RNAi control, with the individual RPG knockdowns having slightly less severe phenotypes than knockdown of either Rib or Trf2 (**Fig. 7C**).

Altogether, the physical interactions of Rib with Trf2, M1BP, and Dref suggest that Rib—either as a subunit of a multi-protein complex, or as a combinatorial interaction partner—cooperates with these known activators of RPG transcription to coordinately increase expression of RPGs. The cell growth deficiency with RNAi knockdown of Trf2— a confirmed RPG transcription initiator—and with RNAi knockdown of individual RPGs, further strengthens the connection between RPG transcription and embryonic SG cell growth.

### Rib-dependent tracheal cell growth is not linked to RPG binding

Finally, we asked whether loss of Rib affects cell growth in other embryonic tubes and, if so, whether the mechanism of Rib action to mediate cell growth is the same as in the embryonic SG, *i.e.*, through boosting RPG transcription. We investigated the embryonic tracheal dorsal trunk (DT), a multicellular tube, which shows enriched Rib expression and exhibits failed tube elongation with *rib* loss (Bradley and Andrew, 2001; Kerman et al., 2008; Shim et al., 2001). Morphological and volumetric analysis of *rib* mutant DT cells revealed comparable cell number (≈ 10 cells per DT segment), but a remarkable decrease in cell growth, with *rib* mutant DT cells achieving only ≈ 46% the size of WT DT cells (**Fig. S6A – B**). ChIP-Seq experiments using two tracheal drivers, *trh*-GAL4 and *btl*-GAL4, to drive expression of *rib*-GFP revealed a relatively large number of Rib-bound tracheal genes—1433 unique genes— that were largely distinct from Rib-bound SG genes (**Fig. S6C, Tables S5**). Importantly, in striking contrast to the SG GO clusters, *cytoplasmic translation/ribosome* did not feature in the top 10 (or even top 92) DAVID GO terms for Rib-bound tracheal genes with only a small fraction of tracheal RPGs bound by Rib, i.e., 13 of 84 (**Table S2**). Notably, the binding of Rib to the majority of tracheal RPGs was not TSS-proximal (**Fig. S6D**). Thus, despite the shared requirement for Rib in embryonic cell growth of both tubular organs, Rib-dependent growth in the trachea is likely through regulation of alternative growth-promoting factors.

## DISCUSSION

The findings described herein reveal that post-mitotic growth is a key feature of embryonic morphogenesis and development. WT SG cells more than double their size as they undergo tube morphogenesis by internalization of the primordia, tube elongation, and collective migration. The nuclear BTB factor Rib plays a major role in post-mitotic early cell growth in at least two epithelial tubular organs. Rib coordinates growth in the two tissues by apparently different mechanisms. In the SG, Rib appears to mediate growth by boosting translational capacity; Rib binds the TSS of nearly every SG-expressed RPG (**Fig. 4, S1A – S1E**) as well as key components of the translational machinery (**Fig. S7A)**. Moreover, Rib is required for full expression of every RPG tested, presumably acting through its association with both known and yet-to-be-discovered sequence-specific activators of RPG expression (Baumann and Gilmour, 2017; Yamashita et al., 2007), as well as through Trf2, the transcription initiator of RPGs (Wang et al., 2014). Despite *rib* loss impacting post-mitotic growth of the embryonic trachea, Rib does not bind the majority of RPGs in the trachea, suggesting that it regulates growth through other targets in this tissue.

### The RPG regulatory program

Although RPGs are closely conserved across species, components of their transcriptional regulatory programs, *i.e.*, sequence motifs and transcription regulators, are highly divergent (Ma et al., 2009). The fast-evolving RPG regulatory programs incorporate layers of redundancies and novelties in higher organisms, and, therefore, elude their abstract portrayal as one universal, cross-species RPG gene regulatory network (Perry, 2005; Tanay et al., 2005). Despite the divergence of RPG regulatory elements between yeast (the model where RPG regulation is relatively well characterized) and mammals, some degree of conservation prevails between *Drosophila* and mammalian (including human) components of RPG regulation in the form of the T-rich sequence (immunoprecipitated by Rib), which is in apposition with the TSS of individual RPGs (Vo Ngoc et al., 2019). The ubiquity of sequence-specific transcription factor binding sites embedded within the promoter-proximal enhancers of RPGs, furthermore, is the least common denominator of conservation within RPG regulatory networks. It is also an indication of the importance of these factors—Rib, Dref, and M1BP in *Drosophila*; GABP, Sp1, and YY1 in mammals—in directing the expression of RPGs as part of a redundant or context-specific transcription program that could be tapped for specialized cell functions requiring augmented translation, e.g., secretion. The function of Btbd18, the predicted mammalian Rib orthologue (FlyBase DIOPT v8.0), in RPG regulation is unclear although its high expression in adult mouse testis parallels that of Rib’s expression in *Drosophila* (Zhou et al., 2017). That the factors—Rib, Dref, and M1BP—may indeed, provide context-dependent regulation is demonstrated by their multifarious roles, including, in the transcription of non-RPGs during embryogenesis (Hirose et al., 1996; Li and Gilmour, 2013; Loganathan et al., 2016). To our knowledge, Rib, a known morphogenetic regulator during embryogenesis, is also the first known factor that interacts with all three of the RPG regulators (M1BP, Dref, and Trf2) and binds all RPGs. That Rib function is critical for full levels of expression of all RPGs and for cell volume gain reveals a SG secretory cell growth program realized by coordinate transcriptional upregulation of RPGs.

### Post-mitotic early cell growth

Despite the importance of growth for tissue structure and function—*e.g.*, oogenesis, muscle hypertrophy, synaptic potentiation—information regarding this process, particularly during embryogenesis, is scarce. However, post-embryonic cell/tissue growth, also referred to as “late growth,” has been recognized as a major developmental strategy in metazoans (O’Farrell, 2004). Late growth in *Drosophila* is best defined for the larval stage, and occurs in two major forms: (i) hypertrophic growth in all differentiated tissues—with the exception of the nervous system—by DNA polyteny (Orr-Weaver, 2015), and (ii) proliferative growth in imaginal tissues—the precursors of adult structures (Irvine and Harvey, 2015; Peng et al., 2009). These processes enable the nearly 1000-fold volume increase that occurs during larval stages and they prefigure the larval-to-adult tissue-mass conversion that occurs—within the pupa—by histolysis of larval tissues and concurrent morphogenesis of adult structures from imaginal cells. Early growth, *i.e.*, embryonic cell/tissue growth, however, has been ascribed exclusively to the cell-growth-free proliferation resulting from early mitotic cycles (O’Farrell, 2004). Thus, studies on the role of cell growth (size or volume gain), if any, during organogenesis in the embryo has been lacking. Analogous to the late (larval) cell growth that prefigures adult organ structure and function, early (embryonic) cell growth that primes larval organs for specialized functions is not implausible. Bolstering this hypothesis is the evidence for early endocycles in several organs—including the single SG endocycle—during their assembly in the embryo (Smith and Orr-Weaver, 1991). Indeed, early growth in the absence of post-blastoderm cell divisions has been noted in the embryonic cells of the nervous system, although the underlying mechanism is unknown (Hartenstein and Posakony, 1990). Thus, early growth of tissues during post-mitotic organogenesis may be a requirement for priming organ function immediately ensuing embryogenesis.

Both RPs and rRNAs are associated with cellular late growth in *Drosophila* via the characteristic “Minute” mutants (each *Minute* mutation results from heterozygosity for the loss of function of a single RPG) and the “*diminutive* (*dm*)” mutant (loss-of-function of *dMyc*, a key activator of rRNA expression); both *Minute* and *dm* mutations are associated with decreased cell size, decreased body size, decreased ability to compete with neighboring WT cells, and decreased viability (Baker, 2020; Grewal et al., 2005; Johnston et al., 1999; Marygold et al., 2007; Morata and Ripoll, 1975). Our analysis demonstrates that cellular early growth does occur in the embryonic SG, dovetailed with tube morphogenesis. Indeed, both SG and tracheal cell growth are impaired in the absence of Rib. Thus, our previous finding of significant decreases in apical membrane expansion of *rib* mutant SG and trachea with incomplete tube elongation could be either linked to this cell growth deficiency and/or to the altered expression of Rib-dependent morphogenetic regulators (Kerman et al., 2008). Hence, early cell growth may constitute yet another morphogenetic strategy for epithelial tube elongation in addition to the other known strategies of oriented cell division (Baena-Lopez et al., 2005; Concha and Adams, 1998; Saburi et al., 2008), cell rearrangement/intercalation (Blankenship et al., 2006; Keller, 1980; Saxena et al., 2014), and cell shape changes (Diaz-de-la-Loza et al., 2018; Paluch and Heisenberg, 2009).

The tissue-specific context for RPG regulation by Rib during SG tubulogenesis is likely a manifestation of the gland’s primary function: high-level protein secretion. It is possible, hence, that the RPG regulation by Rib accommodates, in addition to the early growth of SG secretory cells, their sustained requirement to boost secretory output. Therefore, the Rib-dependent cell growth program might be intertwined with the CrebA-dependent cell secretion program (Fox et al., 2010) as both require high fidelity translational machinery. CrebA, a bZIP transcription factor, upregulates the secretory pathway component genes over the timeframe that is concordant with Rib-mediated early cell growth (Johnson et al., 2020), and one of the CrebA orthologues has been implicated in also scaling the translational capacity of hormone-secreting mammalian cells (Khetchoumian et al., 2019). The temporal dynamics of transcript expression for all the major factors implicated in this hypothetical gene regulatory network, viz., Rib (RPGs), Trf2 (RPGs), M1BP (RPGs), Dref (RPGs), Myc (rRNAs), and CrebA (secretome), moreover, are strikingly similar (**Fig. 7A**). Of particular note is the very high-level expression of Myc, which is known to upregulate ribosomal RNAs (**S7B – C**). Hence, a parsimonious working model in which the RPG regulatory program is bipotent in meeting the translation requirements for both cell growth and secretion could be postulated based both on our present results and other previously published studies. The general applicability/possibility, moreover, of targeted, *i.e.*, tissue-specific, early growth programs as a developmental strategy during embryogenesis for other organs undergoing eutelic (post-mitotic) morphogenesis remains to be tested; nonetheless, our tracheal-specific Rib ChIP-Seq findings and volumetry certainly lend support to this hypothesis.

In summary, determination of the SG-specific direct transcriptional targets of Rib revealed the ribosome as the primary target, implicating the translational machinery in mediating post-mitotic early cell growth that was neither previously defined nor considered relevant for organ formation in *Drosophila*. Embryonic SG tubulogenesis, thus, provides a cogent demonstration of early cell growth melded into an organ (tube) morphogenetic program. Due to the principal secretory role of the SG, we postulate that early cell growth is likely an ineluctable accommodation resulting from the augmented translation capacity required for the mobilization of the SG secretome.

Although ribbon (*rib*) was named for defects in the mutant larval denticle structures (Nusslein-Volhard et al., 1984), the choice for its name—in light of evidence for its role as a customized regulator of the ribosome—is quite prescient.

## SUPPLEMENTAL INFORMATION

Supplementary information includes seven figures and five tables.

## ACKNOWLEDGMENTS

The authors thank M. Chiu, K. Kim, S. Lannon, and M. Luchetti for ranking FISH images; C. Talbot for assistance with the volcano plot; S. Celniker for sharing the RNA-seq data of SG-expressed RPGs; S. Beckendorf, J. Kadonaga, D. Gilmour, M. Yamaguchi, M. Siomi, D. Cavener, and K. White for sharing antibodies; C. Nusslein-Volhard, M. Krasnow, and S. Hayashi for sharing fly lines; E. Chen for sharing the S2R+ cell line; D. Barrick, J. Berger, and G. Seydoux for their comments to help improve an earlier version of the manuscript; FlyBase, BDGP, BDSC, VDRC, and DSHB for curation and distribution of valuable reagents and resources for the fly community; staff from the JHMI SOM microscopy core and the transcriptomics and deep sequencing core for assistance. This work was supported by NIH grants R01DE013899 (D.J.A) and R35GM119553 (M.S.).

## AUTHOR CONTRIBUTIONS

R.L. and D.J.A. conceived the project and designed the experiments; R.L., H.C., Y.W., and D.J.A. performed *Drosophila* embryo experiments; R.L. did the confocal imaging; R.L. performed the EMSAs; J.H.K. performed S2R+ cell culture; R.L. performed the Co-IPs with guidance from J.H.K.; M.S. performed the ChIP-Seq; D. C. L. and M.B.W. performed ChIP-Seq data analysis with guidance from M.S.; M.B.W. performed the RT-qPCR; R.L. and D. C. L. performed DAVID analysis; D.J.A. performed TEM and MEME analysis; R.L. and D.J.A. wrote the manuscript with input from all the authors; D.J.A. supervised the project.

## SUPPLEMENTARY INFORMATION

**Figure S1.**
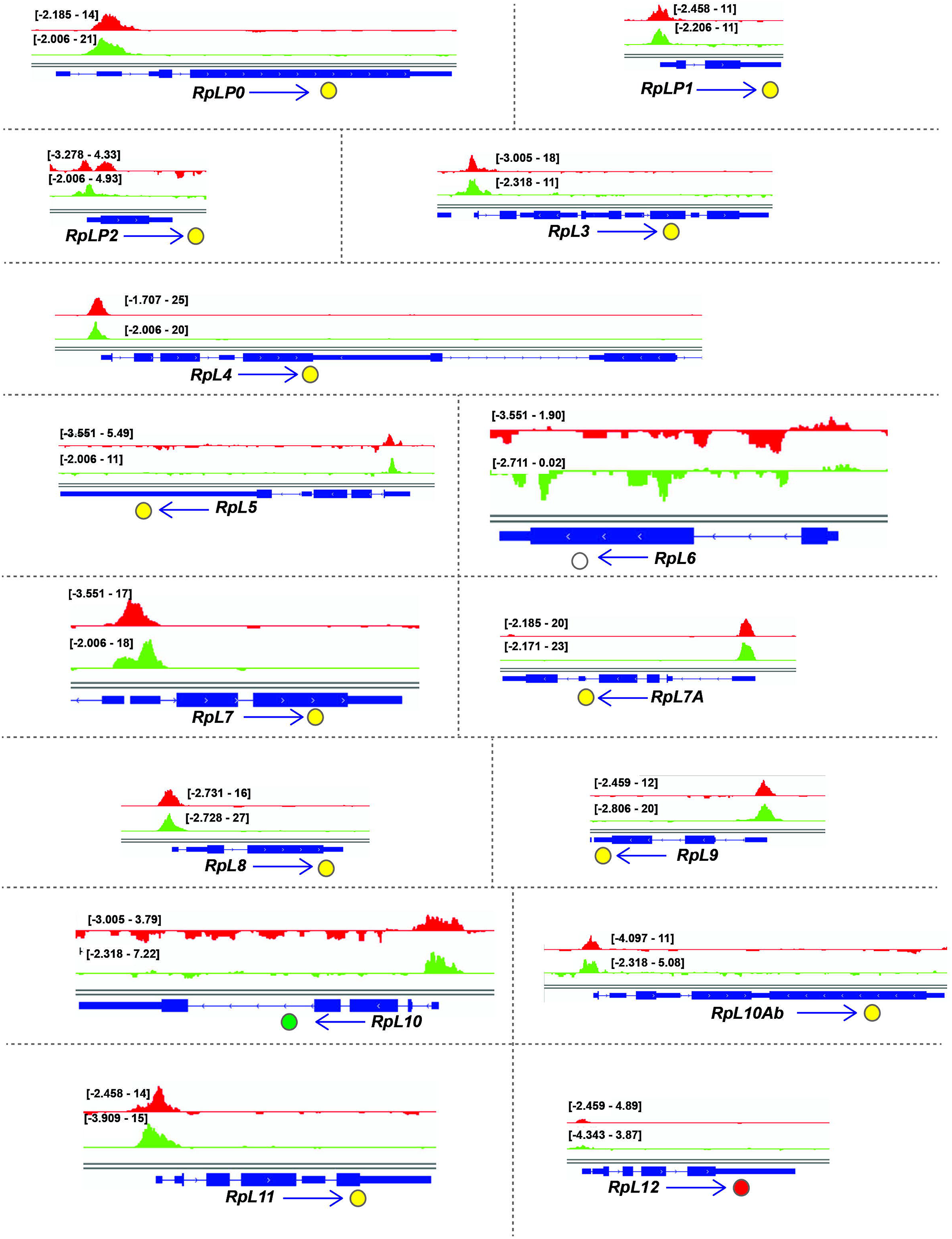
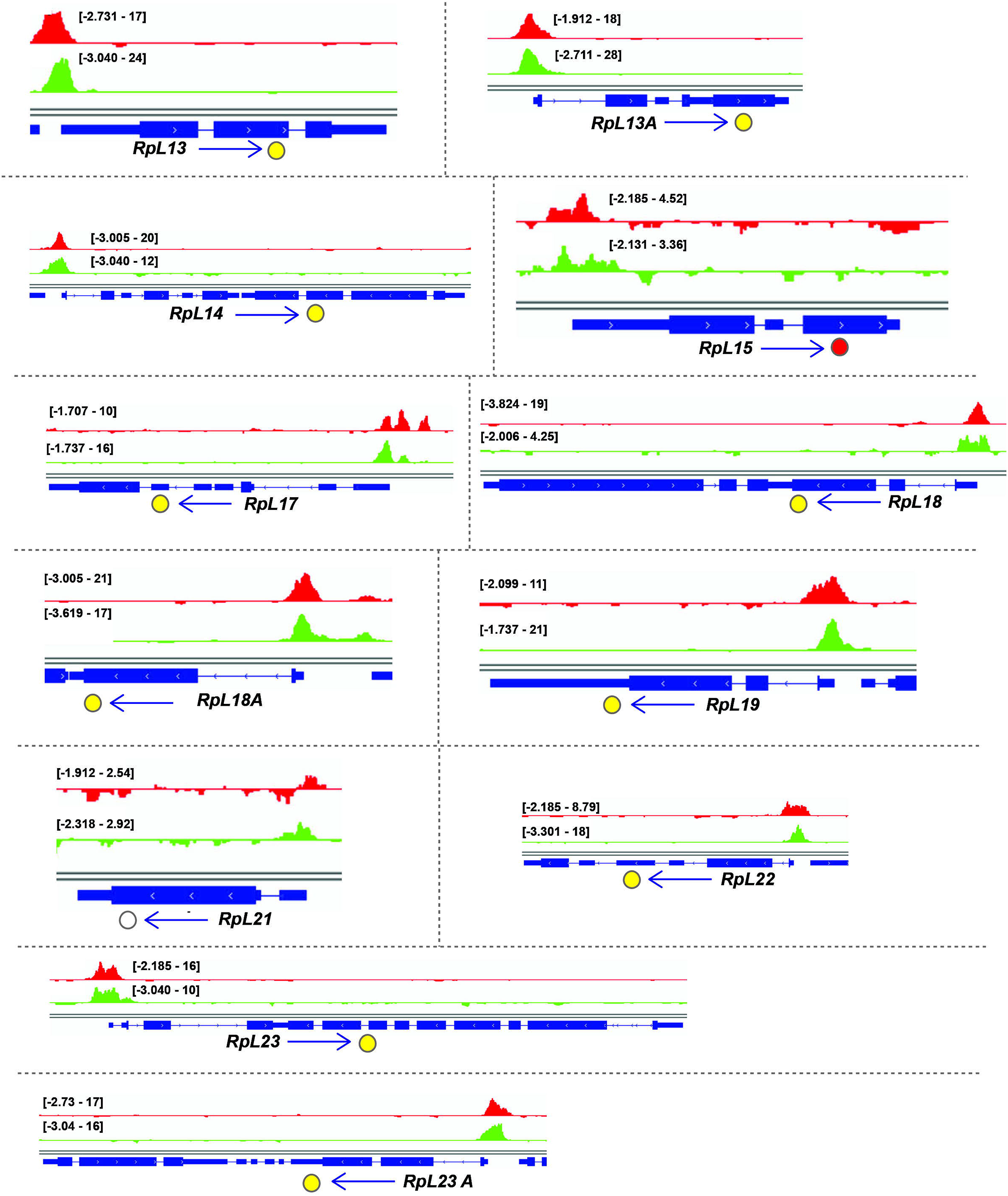
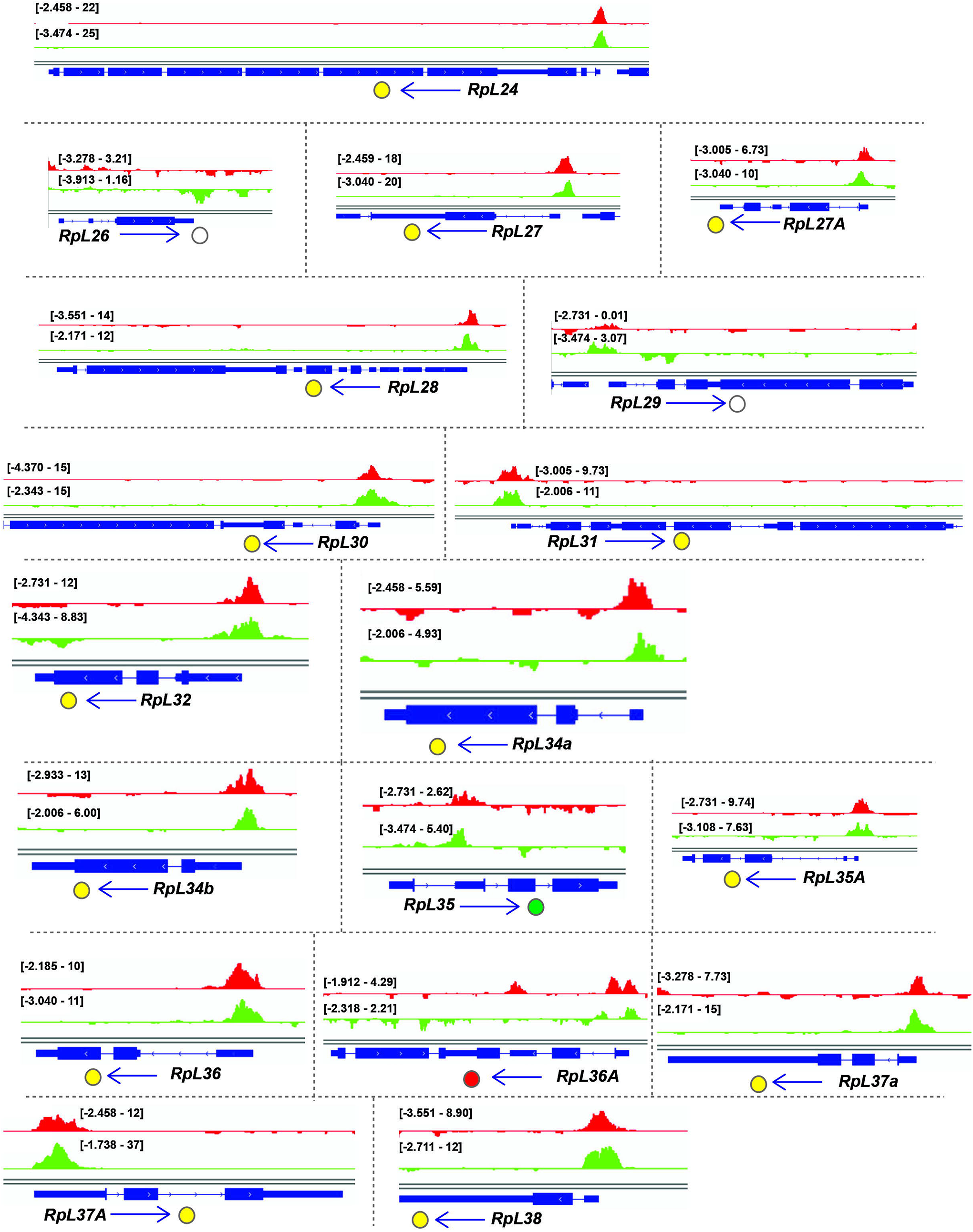
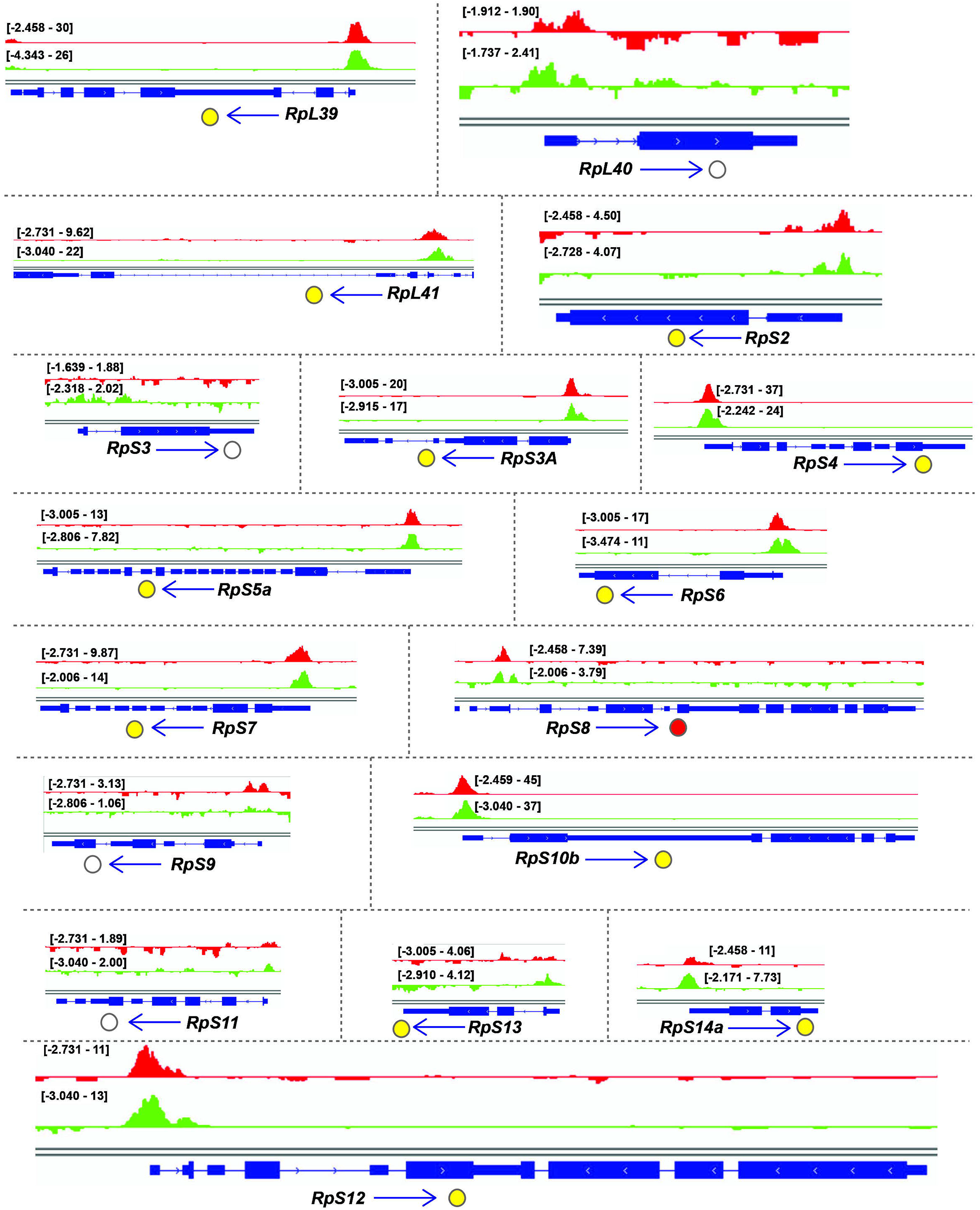
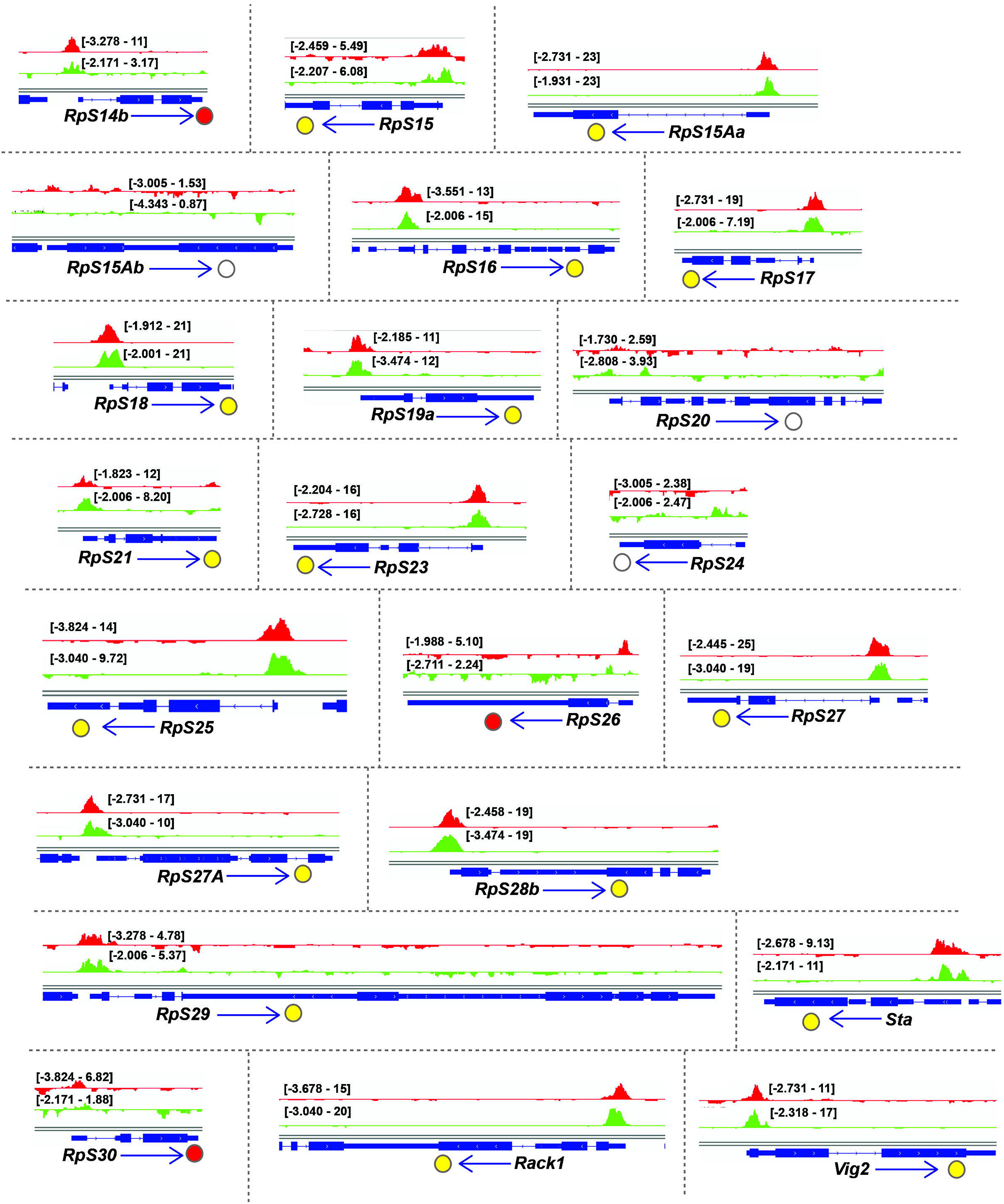
Binding tracks from Rib-GFP ChIP-Seq for all SG-expressed RPGs. *(related to* Fig. 4) (**A – E**) *fkh*-GAL4 > UAS-*rib-GFP* tracks are in red. *sage*-GAL4 > UAS-*rib-GFP* tracks are in green. The colored circles correspond to the individual track colors when either of the binding peaks reach the enrichment threshold of log_10_ binding likelihood ≥ 4. In cases where both the binding peaks meet the enrichment threshold, the circles are colored yellow. In cases where both peak thresholds are < 4, the circles are unfilled. Note that even when the enrichment threshold was <4, binding signals for the region shown are often highest at the RPG transcription start site.

**Figure S2.**
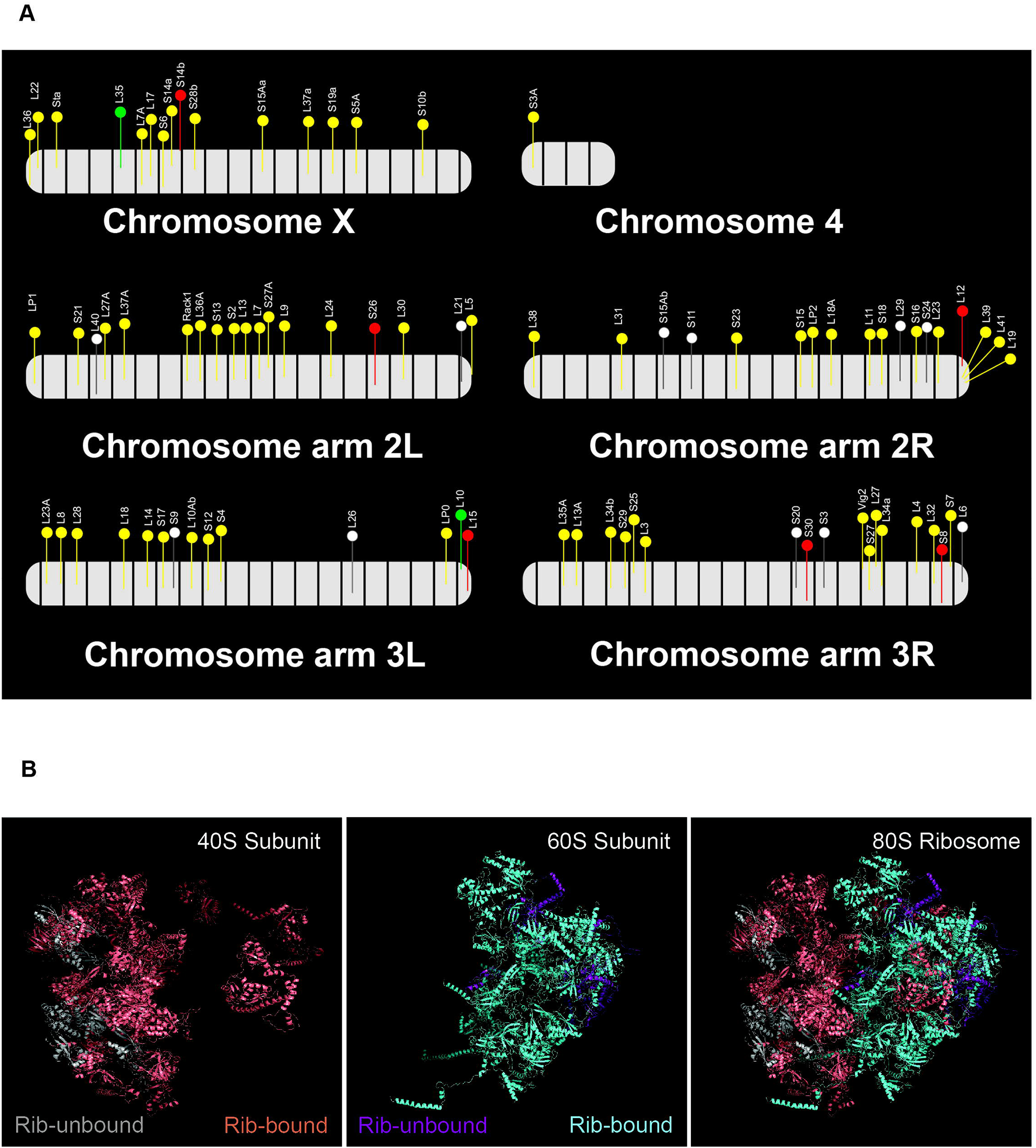
Rib binding of RPGs across the *Drosophila* genome and the position of the corresponding proteins on the ribosomal subunit structures. *(related to* Fig. 4) **(A)** *Drosophila* RPG loci are dispersed across all chromosomes. Colored flags represent RPGs bound by Rib (Log_10_ binding likelihood threshold ≥ 4). Unfilled flags represent genes not bound by Ribbon (Log_10_ binding likelihood threshold < 4). Relative gene positions are approximate representations based on the standard cytogenetic map (FlyBase, FB2017_04). Yellow—genes bound by Rib in both *fkh*-GAL4 and *sage*-GAL4 driver tracks, Red—genes bound by Rib in the *fkh*-GAL4 track only, Green—genes bound by Rib in the *sage*-GAL4 track only. **(B)** Rib binds to genes encoding for proteins of both the 40S and the 60S ribosomal subunits. The proteins for which the corresponding genes lack Rib binding in the ChIP-Seq are shown in gray for the 40S subunit and in purple for the 60S subunit.

**Figure S3.**
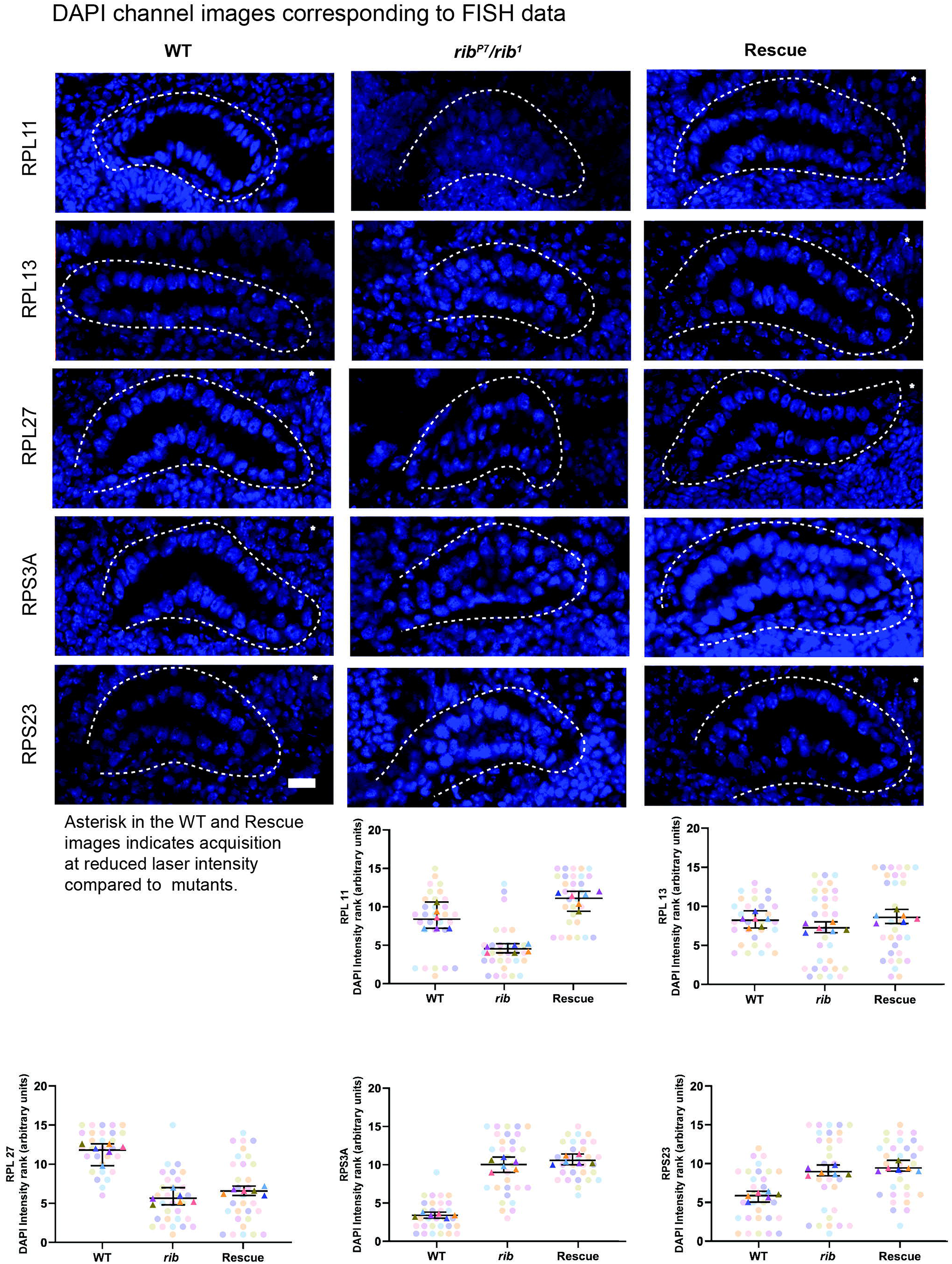
DAPI channel images from the FISH experiment. *(related to* Fig. 5) <u>Top:</u> DAPI channel images for the corresponding representative images from RPG FISH experiments shown in Fig. 5. <u>Bottom:</u> The SuperPlots for the DAPI intensity showed that the ranking of blind samples does not follow an order consistent with the ordering of RPG probe intensity, *i.e.*, WT or *rib* SG rescue followed by the *rib* mutant for RPG probe rankings (Fig. 5) *vs.* no concordant group order for DAPI rankings. Scale bar: 10 µm.

**Figure S4.**
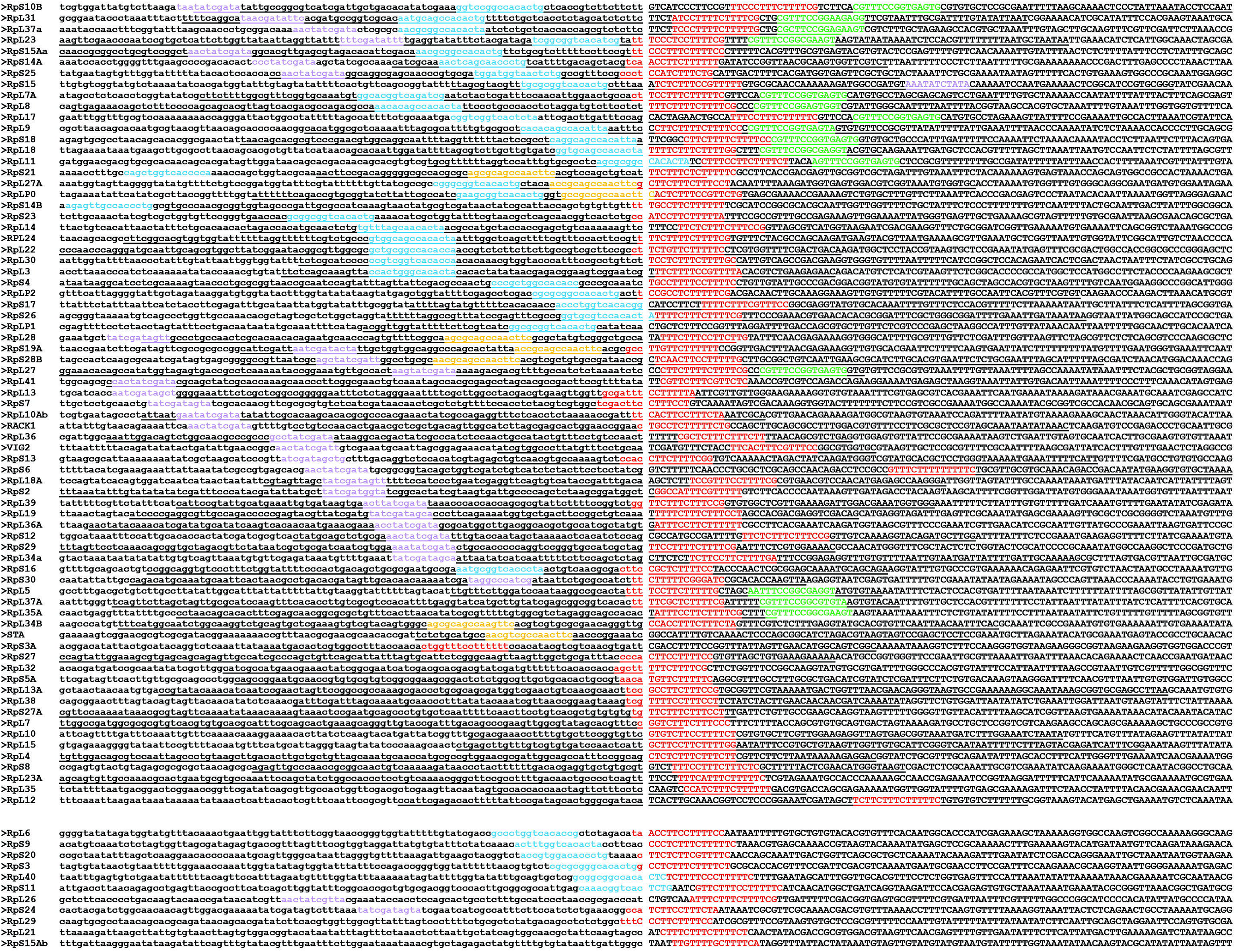
Putative Rib-bound RPG enhancer motifs. *(related to* Fig. 6) Ordering of RPG sequences with putative enhancer motifs from MEME-ChIP to show their positions relative to the transcription start (as determined from Flybase data and indicated by a gap in sequences), from −100 bp to +100 bp. Bases in lower case are upstream of the TSS. Conserved sequence motifs (indicated by colored text and corresponding to Fig. 6A and 6B are observed in genes bound by Rib (upper block) as well as in genes not bound by Rib (lower block). Underlined sequences in the upper block were immunoprecipitated by Rib-GFP in the ChIP-Seq experiments.

**Figure S5.**
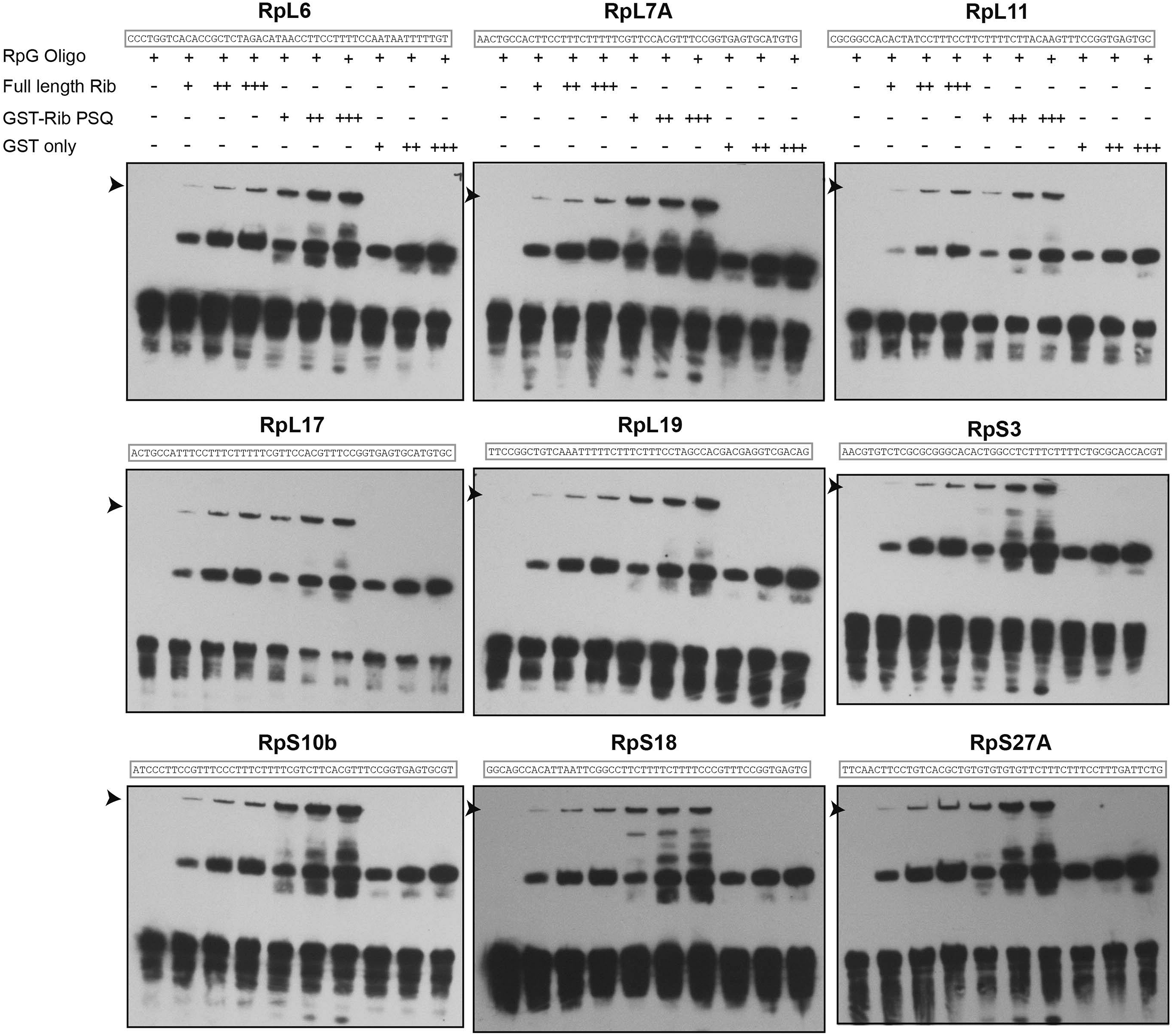
RPG enhancer EMSAs with Rib and Rib DNA-binding domain. *(related to* Fig. 6) RPG enhancer fragments corresponding to the sequences of RpL6, RpL7A, RpL11, RpL17, RpL19, RpS3, RpS10b, RpS18, and RpS27A revealed direct binding by both the full-length Rib protein and Rib-PSQ DNA binding domain. Arrowheads mark Rib-dependent mobility shifts. All EMSAs were performed twice with identical results.

**Figure S6:**
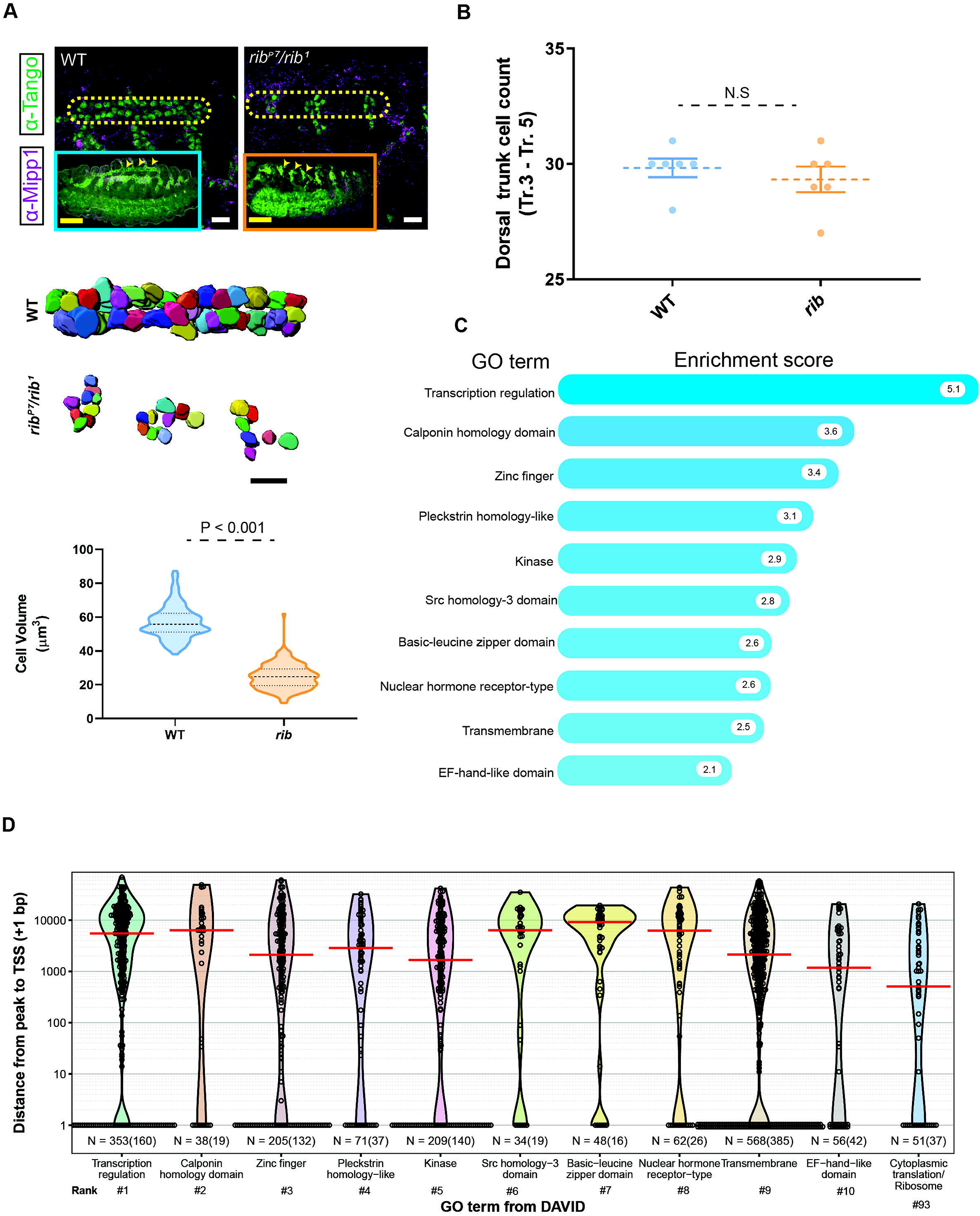
Tracheal DT cell growth deficit in *rib* mutants and Rib-bound tracheal genes. **(A)** <u>Top:</u> Comparison of WT and *rib* mutant trachea from stage 16 embryos. Yellow outline encloses dorsal trunk segments corresponding to Tr.3 – Tr.5 and reveals the failure in tube elongation and fusion in *rib* mutants. Scale bar: 10 µm. <u>Inset:</u> Arrowheads indicate dorsal trunk segments from Tr.3 – Tr.5 in the whole embryo. Scale bar: 50 µm. <u>Bottom:</u> Tracheal cell volumetry shows decreased size in *rib* mutant dorsal trunk cells compared to those from stage-matched wild-type embryos. A total of 260 cells (5 embryos/group; stages 15, 16) with clear marker staining, from segments Tr.3 – Tr.5, whose entire volume could be unambiguously measured, were volume-rendered (two-tailed, unpaired t-test; group median and quartiles labeled). Scale bar: 10 µm. **(B)** Cell counts of WT and *rib* mutant tracheal dorsal trunk (Tr. 3 – Tr. 5). (n=6 embryos/group; two-tailed, unpaired t-test; mean with SEM). N.S.—Not significant. **(C)** Functional clustering of Rib-bound tracheal genes under gene ontology categories (GO terms) according to the Database for Annotation, Visualization and Integrated Discovery (DAVID). The top ten GO terms with their associated enrichment scores are shown. *See* ***Table S5*** *for meta data*. **(D)** Mapping of Rib binding peaks relative to the TSS of the closest gene and sorted by DAVID GO terms in the embryonic trachea shows a preponderance of target binding peaks localizing to regions distant from the TSS compared to the SG (Fig. 3D). Rib binds only a small fraction of RPGs (GO cluster rank #93), and primarily to sequences distal from the TSS-proximal promoter. Red line indicates median distance from the peak to TSS (+1 bp). N = number of peaks versus (number of genes).

**Figure S7:**
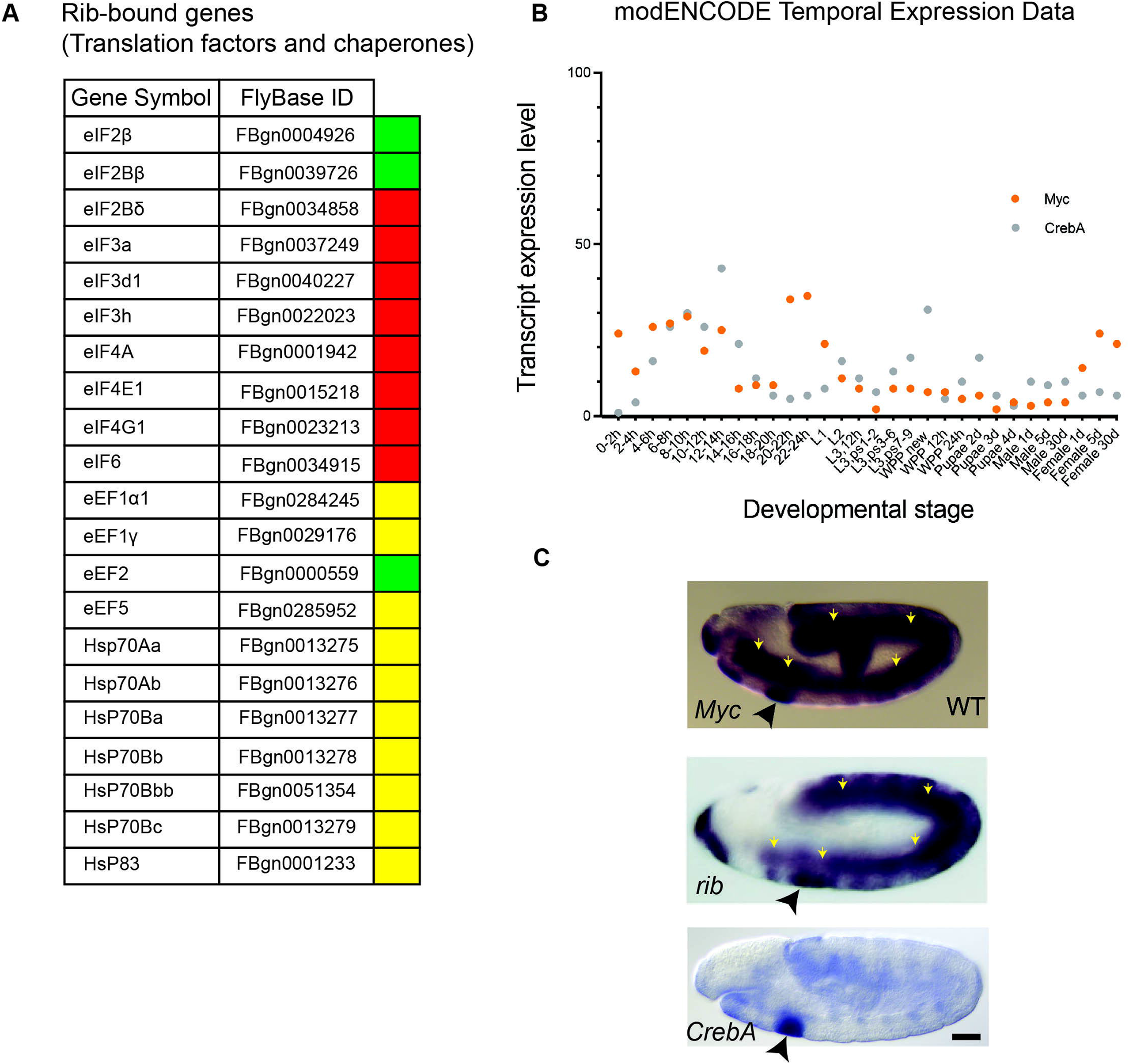
Rib binds translation factors and chaperones for protein synthesis. *(related to discussion)* **(A)** List of Rib-bound translation factors and chaperones from the embryonic SG ChIP-Seq. Genes are color coded according to whether binding peaks for each experiment was ≥4 in either *fkh*-GAL4 track only (red), *sage*-GAL4 track only (green), or both *fkh*-GAL4 and *sage*-GAL4 driver tracks (yellow). **(B)** Transcript expression profiles of *CrebA* and *Myc* during the course of *Drosophila* development correspond closely to the temporal expression profiles of other RPG regulatory factors shown in Fig. 7A. Note the difference in Y-axis range compared to Fig. 7A. **(C)** In situs from early stage embryos with probes for *dm (Dmyc), rib*, and *CrebA,* reveal high-level SG expression (black arrowhead) with all three genes. Note also high-level expression of *dm* and *rib*, but not *CrebA*, in the mesoderm (yellow arrowheads). Scale bar: 50 µm.

**Table S1.** (related to **Fig. 3**) List of the top ten annotation clusters with associated enrichment scores and P-Values from DAVID analysis of genes bound by Rib in the SG.

**Table S2.** (related to **Fig. 4**) List of RPGs expressed in third instar larval (L3) and white prepupal (WPP) SGs showing values from Rib-binding data from embryonic tissue-specific ChIP-Seq (yes or no, for above or below binding likelihood threshold), their fold changes in *rib* versus WT whole embryonic RNA microarray analyses, as well as the overall transcript levels in the SG from BDGP-generated RNA-seq data (kindly provided by Sue Celnicker).

**Table S3.** (related to **Fig. 4**) List of primers used for RT-qPCR experiments.

**Table S4. (**related to **Fig. 6**) List of oligos used for EMSA experiments.

**Table S5.** (related to **Fig. S6**) List of the top ten annotation clusters with associated enrichment scores and P-Values from DAVID analysis of genes bound by Rib in the Trachea.

## MATERIALS AND METHODS

### Fly Strains

Oregon R (BDSC stock number 2376) embryos were the wild-type control in all experiments. The trans-allelic combination of *rib^1^* (stock number 3240) (Bradley and Andrew, 2001) and *rib^P7^* (Shim et al., 2001) was used for *rib* mutant phenotype analysis, cell volumetry, gene expression analysis by microarray, qRT-PCR and *in situ* hybridization. These alleles are EMS-induced mutations generated at different times and in different labs; *rib^P7^* has a premature stop codon at residue 22 and *rib^1^* has a premature stop codon at residue 283. *UAS-rib-GFP* was built by cloning a PCR amplification of the full-length *rib* ORF into the pENTR-D vector and subsequent gateway cloning into the pTWG vector, placing the entire GFP coding region downstream of and in frame with the Rib ORF. The following lines were generated to test for rescue of *rib* mutant SG phenotype*: rib^1^ fkh-GAL4/CyO, ftz-lacZ and rib^P7^ UAS-rib-GFP/Cyo, ftz-lacZ.* UAS-Nuclear-lacZ expression driven by *fkh-GAL4, sage-GAL4* and *trh-GAL4* was used to determine the full set of cell types where these drivers are active. The RNAi line for *rib* was procured from VDRC (stock number 103977). RNAi lines for Trf2, RpL19 and RpS29 were obtained from BDSC (stock numbers 36835, 65117, and 67889 respectively). SG-specific RNAi was driven by *fkh-Gal4; UAS-Dicer2.* The *trh*-GAL4 construct was made by PCR amplifying the *trh* enhancer region using primers (**Table S3B**) for integration into the pChs-Gal4 multiple cloning site by restriction digest with SacII and BamH1 (Sotillos et al., 2010).

### Drosophila Husbandry

Flies were raised at 25°C on a standard yeast/molasses medium (1212.5 mL water, 14.7 mL agar, 20.4 g yeast, 81.8 g cornmeal, 109.1 mL molasses, 10.9 mL Tegosept, 3.4 mL propionic acid, 0.4 mL phosphoric acid). All experiments were conducted on embryos collected at 25°C except for those used in the RNAi knockdown experiments, which were collected at 28°C.

### Antibody production and Immunostaining

Antiserum for Rib was generated in guinea pig to the product of the full-length *rib* ORF subcloned into the BamH I site of pET-15b vector (Novagen). Recombinant full-length Rib was expressed and purified from *E. Coli* as inclusion bodies and injected into the host animal following standard immunization protocols (Covance).

For immunostaining, embryos were fixed in 4% formaldehyde (Sigma; Cat#252549) in PBS with an equal volume of heptane (Sigma; Cat#34873) for 25 minutes at room temperature, rinsed in MeOH, washed several times in PBS containing 0.1% Triton-X-100 (PBT), and incubated overnight with primary antibodies at 4 °C. The next day, embryos were washed, blocked, and incubated with secondary antibodies at 1/200 dilution for 2 hours at room temperature.

After washing, embryos were counterstained with DAPI (1:1000), and mounted in Aqua-Polymount (Polysciences; Cat#18606-100). For Avidin and Biotin-HRP based staining reactions, vectastain ABC kit (Vectastain Laboratories; Cat# PK-4001) was used.

Antisera were used at the following final concentrations:

Rat anti-Ribbon (1:50); Rabit anti-SAS (1:1000); Guinea Pig anti-Ribbon (1:1000); Guinea Pig anti-Mipp1 (1:500); Guinea Pig anti-Sage (1:100); Rabbit anti-CrebA (1:5000); Rat anti-CrebA (1:1000); Rat anti-Dead ringer (1:5000); Rabbit anti-Fork head, a gift from S. Beckendorf (1:500); Mouse anti-DCP1, a gift from M. Siomi (1:1000); Rabbit anti-PKC ζ (RRID: AB_2300359) (1:500); Mouse anti-Fibrillarin (RRID: AB_523649) (38F3) (1:200); Rabbit anti-GFP (RRID: 221569) (1:1000); Mouse anti-Tango (RRID: AB_528486) (1:2); and Rabbit anti-βGalactosidase (RRID: AB_221539) (1:500).

The following secondary antibodies were used:

Biotin-conjugated goat anti-mouse IgG (H+L) (RRID: AB_2533969); Biotin-conjugated goat anti-rabbit IgG (H+L) (RRID: AB_2533969); Biotin-conjugated goat anti-rat IgG (H+L) (RRID:AB_2535646); Alexa Fluor 488-conjugated goat anti-mouse IgG (RRID: AB_2633275); Alexa Fluor 488-conjugated goat anti-rabbit IgG (RRID: AB_2633280);

Alexa Fluor 488-conjugated goat anti-rat IgG (RRID: AB_2534074); Alexa Fluor 568-conjugated goat anti-rabbit IgG (RRID: AB_143157); Alexa Fluor 568-conjugated goat anti-guinea pig IgG (RRID: AB_141954); Alexa Fluor 647-conjugated goat anti-rabbit IgG (RRID: AB_2633282); and Alexa Fluor 647-conjugated goat anti-guinea pig IgG (RRID: AB_141882).

### Transmission Electron Microscopy

Embryos were processed for TEM by first dechorionating them in 50% bleach and then fixing them in 5% glutaraldehyde and heptane. After manual devitellinization, embryos were fixed in 4% glutaraldehyde (Polysciences; Cat#18428-5) and 2% acrolein (Sigma; Cat#110221) in 0.1 M cacodylate buffer (Sigma; Cat#97068). Devitellinized embryos were transferred to a chilled mixture of 1% osmium tetroxide (Polysciences; Cat#23311-10) and 2% glutaraldehyde in 0.1 M cacodylate buffer, and then post-fixed in 1% osmium tetroxide in 0.1 M cacodylate buffer. Fixed embryos were dehydrated and embedded in Epon as previously described (Myat and Andrew, 2000a). Sections, obtained on a Reichart-Jung Ultracut E, were stained with 2% uranyl acetate (Polysciences; Cat#6159-44-0) and lead citrate (Polysciences; Cat#25350-100) for viewing.

### Chromatin immunoprecipitation and deep sequencing (ChIP-Seq)

Chromatin extraction and immunoprecipitation were performed as described previously (Negre et al., 2006). Briefly, chromatin from multiple independent collections of stage 11 – 16 *fkh*-GAL4::UAS-*rib-GFP*, *sage*-GAL4::UAS-*rib-GFP, btl-GAL4::UAS-rib-GFP and trh-GAL4::UAS-rib-GFP* embryos were cross-linked at room temperature in 1.8% formaldehyde in 2 ml of homogenization buffer (60 mM KCl, 15 mM NaCl, 15 mM HEPES {pH7.6}, 4 mM MgCl2, 0.5 mM DTT, 0.5% Triton X-100 and cOmplete™ protease inhibitor cocktail [1 tablet per 50 ml buffer]). The cross-linked material was resuspended in 0.1%SDS and 0.5% N-lauroylsarcosine in 0.5 ml lysis buffer (140mM NaCl, 15mM HEPES [pH 7.6], 1 mM EDTA, 0.5 mM EGTA, 0.1% sodium deoxycholate, 1% Triton X-100, 0.5mM DTT and cOmplete™ protease inhibitor cocktail [1 tablet per 50 ml buffer]). Chromatin was sonicated three times at 4°C using the Sonic Dismembrator Model 100 (Fisher Scientific) under the following conditions: power setting 3, 20s ON, 20s OFF. Immediately after sonication, the chromatin extract was stored at −80°C prior to immunoprecipitation. Immunoprecipitations were performed, as described in (Negre et al., 2006), using a polyclonal goat anti-GFP antibody, gift from K. White (1:250). Immunoprecipitated DNA was prepared for Illumina sequencing using the Illumina TruSeq ChIP Sample Prep Kit (*fkh*-GAL4::UAS-*rib-GFP* and *sage*-GAL4::UAS-*rib-GFP* samples), the NuGen Ovation Ultralow Library System (*btl-*GAL4*::UAS-rib-GFP* and *trh-*GAL4*::UAS-rib-GFP*), or the KAPA DNA HyperPrep Library Prep Kit (independent replicate of *btl-*GAL4*::UAS-rib-GFP*). Illumina sequencing was performed according to the manufacturers specifications at the University of Minnesota Genomics Center (SG and Tr. dataset) or at the UCLA Technology Center for Genomics & Bioinformatics (Tr. dataset).

### ChIP-Seq data processing

Detailed methods for the *sage-*GAL4; UAS*-rib-*GFP and *fkh-*GAL4; UAS*-rib-*GFP ChIP-seq are available from a previous report (Loganathan et al., 2016). Briefly, ChIP-seq peaks (binding sites) were called by comparing biological replicates to an input DNA control (from non-immunoprecipitated chromatin). Sequenced DNA was processed using FASTQC (RRID:SCR_014583) and FASTQ Groomer (Blankenberg et al., 2010), then mapped to the *Drosophila melanogaster* BDGP release 6 (dm6, August 2014) using Burrows-Wheeler Alignment (BWA) tool with default parameters (dos Santos et al., 2015; Li and Durbin, 2009; St Pierre et al., 2014). Sequencing reads from biological replicates were combined after mapping using Picard (http://broadinstitute.github.io/picard) and the MACS (v2) peak caller was used to identify and score peaks (Zhang et al., 2008). Peak calling was carried out using the following MACS parameters (P-value: 10e-5; mfold: 10, 32), comparing ChIP DNA to matching input control samples. These previously archived data, including peaks and log _10_(likelihood ratio) signal files generated from MACS yielded 494 Rib-bound genes, and were used in **Fig. 4** and **Supplementary Figures (Fig. 1A – Fig.1E)** and are accessible in GEO (GSE73781). Data were displayed in the Integrated Genomics Viewer (Broad Institute; RRID:SCR_011793). In gene browser tracks, the X axis is genomic position, and the Y axis is log _10_-normalized fragments per kilobase mapped, as in (Loganathan et al., 2016).

To obtain a set of more highly reproducible peaks, and thus reduce the noise and/or spurious peak calls from individual ChIP-Seq replicates, the original ChIP-Seq reads for the *UAS-rib-GFP* experiments were re-processed and analyzed using a ChIP-Seq pipeline featuring the more stringent irreproducible discovery rate (IDR) correction (Li et al., 2011). This yielded 436 peaks that mapped to 413 genes with RPGs recovered as the most enriched cluster.

Briefly, the reads were first trimmed using Trimmomatic (Bolger et al., 2014) to remove sequencing adapters and low-quality reads. Mapping, alignment, peak-calling and IDR thresholding (at *p* < 0.1) were then performed using the ENCODE ChIP-seq IDR pipeline (RRID:SCR_017237) (https://github.com/ENCODE-DCC/chip-seq-pipeline2) for both Rib ChIP experiments. These sets of IDR ChIP-Seq peaks for *fkh-* and *sage-*driven Rib::GFP, along with their IDR pipeline-generated pooled replicate fold-enrichment signal from MACS, were used in all other analyses and figures. Overlapping IDR peak regions from the Fkh- and Sage-driven Rib::GFP ChIP experiments were manually inspected against the *Drosophila* dm6 genome using the IGV browser.

For the comparison of ChIP-seq experiments in **Fig. 3A**, the SG Rib peak set was generated using the BEDTools (RRID:SCR_006646) *intersect* command (Quinlan and Hall, 2010) with the *fkh-* and *sage-*driven Rib::GFP ChIP-seq IDR peaks and control regions were generated using BEDtools *shuffle* command with these peaks. All these peak sets and the fold-enrichment signal bigwig files were loaded into R with the *rtracklayer* package (Lawrence et al., 2009). The *plyranges* package (Lee et al., 2019) was then used to consolidate and summarize each region by its maximum fold-change signal, all of which were then plotted as hexbins with the *ggplot2* package [https://joss.theoj.org/papers/10.21105/joss.01686].

For the tracheal cell-specific ChIP-Seq, IDR peaks were generated using the same methodology as above with replicate datasets from *trh*-GAL4; UAS-*rib*-GFP and *btl*-GAL4; UAS-*rib*-GFP. Regions recovered from both IDR peak sets were then obtained using BEDTools, and further refined by filtering for overlap with an independent, single replicate *btl-*GAL4; UAS-*rib*-GFP experiment, to recover high confidence binding events similar to those mentioned above for the SG dataset.

### Peak-to-gene association and genomic feature analysis

The SG and tracheal Rib IDR peaks were associated with genes in R (RRID:SCR_001905) by the following method: First, the *Drosophila melanogaster* genome (r6.33) was downloaded from FlyBase (ftp://ftp.flybase.net/genomes/Drosophila_melanogaster/dmel_r6.33_FB2020_02), loaded into R using the *GenomicRanges* and *GenomicFeatures* packages (Lawrence et al., 2013), and filtered to retain only protein-coding genes and their TSS coordinates. The *rtracklayer* and *plyranges* packages were then used to import the various peak files into R ‘link’ them to the nearest protein-coding gene. Peaks which overlapped the TSS of more than one gene were associated with all such overlapped genes. All such Rib-linked genes were used for the subsequent DAVID analysis. For the genomic feature analysis, the *ChIPseeker* (Yu et al., 2015) and *ggplot2* packages were used to calculate and display overlap of peaks with various genomic features of the r6.33 *Drosophila genome*.

A list of all protein-coding genes in which the TSS is within 2kb of an SG Rib peak were recorded and this set was considered for motif analysis. All genes both significantly altered in *rib* by microarray and present in and at two statistical cut-offs, not included in above gene set, were manually inspected for peak-like ChIP signals over log_10_ (likelihood ratio) ≥ 4. Those genes with such a peak-like signal were added to the Rib targets gene set.

### MEME Analysis

Multiple Em for Motif Elicitation (MEME), a bioinformatics analysis tool available within The MEME SUITE (RRID:SCR_001783), was used for RPG enhancer analysis on Rib-bound DNA sequence to identify frequently occurring motifs (Quinlan and Hall, 2010). MEME uses the expectation matrix (Em) algorithm for detection of motifs that have enriched instances in the input sequence sets compared with the genomic background. Among the top five ranking motifs from the Rib-bound sequences for RP genes in MEME, three were known for RP gene regulation and, hence, were investigated further in EMSAs; furthermore, their putative binding factors were tested in Rib Co-IPs.

### DAVID Analysis

Functional clustering of Rib-bound genes from the ChIP-seq data were performed to place them under gene ontological categories (GO terms) according to the Database for Annotation, Visualization and Integrated Discovery (DAVID) version 6.8 (RRID:SCR_001881) (Huang da et al., 2009a, b). The results were then imported into R and the FlyBase gene IDs (FBgn) from each gene ontology cluster were isolated and grouped by gene ontology cluster. The FBgn’s were cross-referenced with the peak-to-gene association lists, and the distance from gene TSS to peak for all genes in each cluster were extracted. These peak-to-TSS distances (plus 1 bp for proper display on the log_10_-transformed Y axis) are plotted by DAVID cluster as beeswarm + violin plots in **Fig. 3** and **Fig. S6** using *ggplot2*.

### Microarray gene expression analysis

Three independent collections of stage 11 – 16 *rib^1^*/*rib^P7^* embryos and three of wild-type embryos were isolated using a COPAS Select large particle FACs (Union Biometrica). Total RNA was isolated by Trizol extraction (ThermoFisher; Cat#15596026) and cleaned up with the Qiagen RNeasy kit (Cat#74004). Total RNA (100 ng) was labeled and amplified using standard Affymetrix protocols. Three samples for each genotype were hybridized to Drosophila Genome 2.0 Chips. Scanned intensity values were normalized using RMA Partek software (RRID: SCR_01186) (Irizarry et al., 2003a; Irizarry et al., 2003b) and statistical analyses were performed using the Spotfire software package (TIBCO; RRID:SCR_008858). Target genes were identified as those that were upregulated/downregulated (1.5 fold-change cutoff, P < 0.05) in *rib^1^*/rib^P7^ embryos when compared with Oregon R controls.

### Quantitative RT-PCR

qRT-PCR validation experiments to confirm microarray gene expression differences observed in WT versus *rib* embryos were conducted as in (Loganathan et al., 2016). Statistical significance was determined by Mann Whitney U tests comparing delta Cq values (Cikos et al., 2007). Data represent four technical and three biological replicates for each gene.

### Fluorescent *In Situ* Hybridization

FISH experiments were performed according to Lécuyer et al., (Lecuyer et al., 2008). Briefly, DIG-labeled antisense probes to RpL11, RpL13, RpL27, RpS3A, and RpS23 were made from the following DGRC clones respectively: LD17235, LD24350, AT27980, LD08549, and GM14585. Embryos were fixed according to the aforementioned method for immunostaining. After a 5 min. wash in PBT, embryos were treated with proteinase K (3 μg/ml) for 13 min. at RT followed by 1h treatment on ice. Proteinase digestion was stopped with glycine wash (2 mg/ml). Embryos were post-fixed for 20 min. in 4% formaldehyde and hybridized at 56 ^0^C O/N. FISH signal was developed with tyramide reaction following O/N incubation of embryos in sheep anti-DIG antibody (Roche; Cat#11222089001) (1:1000) at 4 ^0^C, 1 h treatment at RT with anti-sheep biotin (1:500), and 45 min. treatment with Vectastain AB solution (1:100).

### S2R+ Cell Culture

S2R+ cells (Gift from E. Chen; RRID: CVCL_Z831) were cultured in Schneider’s medium (Gibco; Cat# 21-720-024) supplemented with 10% fetal bovine serum (FBS) (Gibco) and penicillin/streptomycin (Sigma). Effectene (Qiagen; Cat# 301425) was used to transfect cells to express proteins from UAS-*rib* and UAS-*rib*-GFP plasmids driven by Ubiquitin-GAL4 according to the manufacturer’s protocol.

### Co-Immunoprecipitation and Western blot

After harvesting transfected S2R+ cells by centrifugation, they were washed with cold PBS and lysed in Lysis buffer (10 mM Tris, pH 7.4, 150 mM NaCl, 1 mM EDTA, 1% Triton X-100, 0.5% NP-40) containing a protease/phosphatase inhibitor cocktail. After centrifugation, supernatants were incubated with the appropriate antibodies (1:450)—Trf2 or Dref or M1BP—at 4 °C for 2-3 hrs. Protein A/G agarose resin (Pierce; Cat# 20423) was used to precipitate the antibodies. Immunoprecipitated proteins were analyzed by SDS-PAGE. Antisera were used for western blots at the following final concentrations: Rabbit anti-TRF2, gift from J. Kadonaga (1:2000); Rabbit anti-M1BP, gift from D. Gilmour (1:5000); Rabbit anti-DREF, gift from M. Yamaguchi (1:5000); Guinea pig anti-Rib (1:2000); Rabbit anti-GFP (1:5000); HRP-conjugated secondary antibodies were used at the following final concentrations: anti-Guinea pig (1:5000) and anti-rabbit (1:10000).

### Electrophoretic Mobility Shift Assay

The sequence for Rib-PSQ DNA binding domain, same as that used on the previous bacterial one-hybrid experiment (Noyes et al., 2008), was subcloned into BamH I sites of the pGEX-KG vector (ATCC). Recombinant GST-tagged Rib-PSQ and GST alone were expressed in *E. Coli* and the IPTG-induced (1mM) soluble fractions of the lysates were used for EMSAs. Recombinant full-length Rib, which was purified for antiserum production as mentioned above, was also used in the EMSAs. Purified recombinant CrebA was produced as previously reported (Fox et al., 2010). For running EMSAs, plus and minus strand oligonucleotides of 50 bases (for RPGs) and ≈ 30 bases (for P24.1 and TSN-2) were designed and synthesized (Integrated DNA Technologies, Inc.). Both strands were 3′ end-labeled with Biotin-11-UTP using Terminal Deoxynucleotidyl Transferase according to the manufacturer’s protocol. Labeled strands were annealed by heating to 95°C for 5□minutes, and then cooling to RT. Mutant binding site oligos were annealed under the same conditions. Rib-FL and CrebA proteins were expressed and purified in bacteria and their concentration determined by nanodrop spectrophotometer (ThermoFisher Scientific). DNA binding reactions were performed according to LightShift EMSA Optimization and Control Kit protocol (Pierce Biotechnology, ThermoFisher Scientific; Cat# 20148). Binding reactions—using 2μl of GST (4.6 μg/μl), GST-Rib-PSQ (4.6 μg/μl), full-length Rib (4.6 μg/μl), and CrebA (0.5 μg/μl) as starter concentrations with RPG enhancer oligos (5 nM)—were run on a 6% native polyacrylamide gel at 100□V for ≈ 1.5□hours (4°C), transferred O/N onto Biodyne B nylon membranes (ThermoFisher Scientific) at 30□V (4°C), and detected using the chemi-luminescent nucleic acid detection module (Pierce Biotechnology, ThermoFisher Scientific).

### Confocal Imaging

DIC images were obtained on an Axiophot microscope (Zeiss) equipped with ProgRes CapturePro (Jenoptik). Fluorescent images were obtained on an LSM 700 Meta confocal microscope (Zeiss) equipped with Zen software (Zeiss). Electron micrographs were obtained on a Phillips CM120 transmission electron microscope. All images for the FISH experiments were acquired at or below 6% laser intensity per channel, *i.e.*, the threshold at which the probe signals became detectable in the *rib* mutant embryos. For a subset of RPG probes, however, the laser intensity was reduced below the 6% threshold for both the WT and rescue embryos due to their excessive signal intensity.

## QUANTIFICATION AND STATISTICAL ANALYSIS

### 3D Rendering of the Embryonic Salivary Gland and Cell Volumetry

Confocal image sections of embryonic SG cells were segmented and manually volume-rendered using the ‘surfaces’ module of Imaris 7.7.2 (RRID:SCR_007370). Individual cell membrane (or nuclear DNA) boundaries from Z stacks were marked for 3D surface creation and volume measurements. The ‘surfaces’ module of Imaris was also used to 3D-render whole embryos manually and record their volumes.

### Quantification of P body Size

SG P-body area was measured with the volocity 3D image analysis software (RRID:SCR_002668). Maximum intensity projected images were segmented for P-body puncta using the *Find Objects* option. The ROI was manually demarcated by outlining the SG, based on the location of Sage-positive secretory cells, to exclude measurements outside the gland. Offset thresholding on the histogram was adjusted on a sample-to-sample basis to eliminate background staining from being included in the measurements.

### Ranking of FISH Images

Randomly shuffled images were ranked according to their FISH probe intensity levels by six members of the Andrew Lab that were “blind” to the identity of experimental specimens. Five images per experimental group (WT, *rib* mutant, and *rib* SG rescue) were included in the ranking analysis and were scored on a scale of 1 (low signal) through 15 (high signal) according to the staining intensity. Similar scoring was also done on the DAPI-stained images of corresponding SGs. SuperPlots were used for visualization of scores with the result from each observer serving as a “replicate” (Lord et al., 2020) to reveal any orderly or disorderly grouping of ranks between the FISH probe intensity and the corresponding DAPI intensity of specimens in the three experimental groups.

### Graphs and Statistical Analysis

GraphPad Prism software (RRID:SCR_002798) was used to generate graphs, and details of statistical analysis are provided in the figure legends.

## RESOURCE AVAILABILITY

Requests for resources and reagents generated in this study should be addressed to Deborah J. Andrew.

High throughput datasets generated for this study (GSE72598, GSE73781, and GSE181361) are freely accessible at the Gene Expression Omnibus (https://www.ncbi.nlm.nih.gov/geo/).

