## Supplementary material for "The Ribb-osome: Ribbon boosts ribosomal protein gene expression to coordinate organ form and function": Table S1

| GO term | Enrichment Score | PValue | Bonferroni | Benjamini | FDR |
| --- | --- | --- | --- | --- | --- |
| Cytosolic translation/ Ribosome | 27.31 | 3.09E-48 | 3.36E-45 | 3.36E-45 | 3.20E-45 |
| Transcription regulation | 3.87 | 2.67E-06 | 0.003 | 3.22E-04 | 3.06E-04 |
| Cell adhesion | 2.70 | 9.02E-05 | 0.016 | 0.001 | 0.001 |
| Transcription factor | 2.49 | 9.29E-04 | 0.236 | 0.045 | 0.044 |
| Convergent extension | 1.90 | 0.002 | 0.947 | 0.079 | 0.075 |
| Epithelial development | 1.84 | 1.52E-04 | 0.152 | 0.010 | 0.009 |
| Apoptosis | 1.80 | 0.005 | 0.995 | 0.123 | 0.117 |
| Wnt pathway | 1.77 | 0.006 | 0.999 | 0.137 | 0.130 |
| Pleckstrin homology-like | 1.77 | 0.003 | 0.902 | 0.409 | 0.409 |
| Transcriptional repressor/bHLH | 1.66 | 0.002 | 0.314 | 0.019 | 0.017 |

**Supplementary Table 1,** related to Fig. 3

**DAVID analysis of genes bound by Ribbon in the embryonic Salivary Gland – Top 10 annotation clusters**
