## Supplementary material for "The Ribb-osome: Ribbon boosts ribosomal protein gene expression to coordinate organ form and function": Table S2

**Supplementary Table 2,** related to Figure 4.

**Ribosomal protein gene expression data from whole embryo microarray analysis (*rib* mutant vs. WT embryos), and BDGP RNA-Seq data for larval (L3) and white pre-pupal (WPP) salivary gland, with Rib bound genes (within 2kb from the binding peak) in the SG and Trachea.**

| Gene | FlyBase ID | Bound by Rib  (SG) | Bound by Rib  (Tr.) | Microarray fold change  (*rib* vs. WT) | p-val.  (*rib* vs. WT) | BDGP transcript levels  (L3 SG) | BDGP transcript levels (WPP SG) |
| --- | --- | --- | --- | --- | --- | --- | --- |
| RpLP0 | FBgn0000100 | Yes | No | -1.082 | 0.032 | 201 | 456 |
| RpLP1 | FBgn0002593 | Yes | No | 1.012 | 0.129 | 580 | 1630 |
| RpLP2 | FBgn0003274 | Yes | No | -1.120 | 0.03 | 644 | 972 |
| RpL3 | FBgn0020910 | Yes | No | -1.103 | 0.045 | 113 | 583 |
| RpL4 | FBgn0003279 | Yes | Yes | -1.057 | 0.028 | 158 | 552 |
| RpL5 | FBgn0064225 | Yes | No | -1.052 | 0.051 | 270 | 582 |
| RpL6 | FBgn0039857 | No | No | -1.032 | 0.175 | 578 | 2240 |
| RpL7 | FBgn0005593 | Yes | Yes | -1.332 | 0.001 | 97.4 | 327 |
| RpL7A | FBgn0014026 | Yes | Yes | 1.040 | 0.016 | 246 | 611 |
| RpL8 | FBgn0261602 | Yes | Yes | -1.024 | 0.222 | 271 | 507 |
| RpL9 | FBgn0015756 | Yes | No | -1.021 | 0.116 | 149 | 519 |
| RpL10 | FBgn0024733 | Yes | No | -1.018 | 0.215 | 96.2 | 143 |
| RpL10Ab | FBgn0036213 | Yes | No | -1.067 | 0.031 | 250 | 700 |
| RpL11 | FBgn0013325 | Yes | No | -1.051 | 0.104 | 146 | 444 |
| RpL12 | FBgn0034968 | Yes | Yes | -1.281 | 0.021 | 184 | 493 |
| RpL13 | FBgn0011272 | Yes | No | -1.017 | 0.072 | 94.2 | 230 |
| RpL13A | FBgn0037351 | Yes | No | -1.008 | 0.850 | 119 | 213 |
| RpL14 | FBgn0017579 | Yes | No | -1.021 | 0.005 | 164 | 511 |
| RpL15 | FBgn0028697 | Yes | No | -1.029 | 0.038 | 267 | 494 |
| RpL17 | FBgn0029897 | Yes | No | 1.053 | 0.151 | 51.1 | 143 |
| RpL18 | FBgn0035753 | Yes | No | -1.052 | 0.017 | 163 | 416 |
| RpL18A | FBgn0010409 | Yes | No | -1.011 | 0.404 | 220 | 789 |
| RpL19 | FBgn0285950 | Yes | No | -1.038 | 0.097 | 325 | 748 |
| RpL21 | FBgn0032987 | No | No | -1.069 | 0.009 | 1190 | 1430 |
| RpL22 | FBgn0015288 | Yes | No | 1.015 | 0.711 | 91.2 | 262 |
| RpL23 | FBgn0010078 | Yes | No | -1.060 | 0.070 | 335 | 389 |
| RpL23A | FBgn0026372 | Yes | No | -1.021 | 0.045 | 745 | 2530 |
| RpL24 | FBgn0032518 | Yes | Yes | -1.215 | 0.072 | 353 | 574 |
| RpL26 | FBgn0036825 | No | No | 1.042 | 0.040 | 439 | 814 |
| RpL27 | FBgn0039359 | Yes | No | -2.533 | 0.0001 | 447 | 1190 |
| RpL27A | FBgn0285948 | Yes | No | -1.067 | 0.047 | 250 | 735 |
| RpL28 | FBgn0035422 | Yes | No | -2.667 | 0.0004 | 216 | 514 |
| RpL29 | FBgn0016726 | No | No | -1.058 | 0.203 | 168 | 492 |
| RpL30 | FBgn0086710 | Yes | No | -1.155 | 0.032 | 122 | 318 |
| RpL31 | FBgn0285949 | Yes | No | -1.055 | 0.071 | 509 | 1080 |
| RpL32 | FBgn0002626 | Yes | No | -1.058 | 0.04 | 181 | 645 |
| RpL34a | FBgn0039406 | Yes | Yes | -1.233 | 0.007 | 25 | 54.5 |
| RpL34b | FBgn0037686 | Yes | No | -1.139 | 0.042 | 166 | 364 |
| RpL35 | FBgn0029785 | Yes | No | -1.157 | 0.024 | 275 | 381 |
| RpL35A | FBgn0037328 | Yes | No | -1.019 | 0.141 | 51.5 | 92 |
| RpL36 | FBgn0002579 | Yes | No | -1.083 | 0.096 | 391 | 983 |
| RpL36A | FBgn0031980 | Yes | No | -1.108 | 0.005 | 159 | 254 |
| RpL37a | FBgn0030616 | Yes | No | 1.053 | 0.218 | 260 | 676 |
| RpL37A | FBgn0261608 | Yes | No | 1.010 | 0.587 | 128 | 239 |
| RpL38 | FBgn0040007 | Yes | No | -1.231 | 0.040 | 134 | 531 |
| RpL39 | FBgn0023170 | Yes | Yes | -1.047 | 0.339 | 161 | 367 |
| RpL40 | FBgn0003941 | No | No | -1.012 | 0.113 | 722 | 2090 |
| RpL41 | FBgn0066084 | Yes | No | -1.059 | 0.085 | 2040 | 3470 |
| RpS2 | FBgn0004867 | Yes | No | -1.107 | 0.002 | 171 | 255 |
| RpS3 | FBgn0002622 | No | No | 1.007 | 0.723 | 374 | 1200 |
| RpS3A | FBgn0017545 | Yes | Yes | -1.098 | 0.021 | 300 | 379 |
| RpS4 | FBgn0011284 | Yes | No | 1.080 | 0.010 | 42.1 | 198 |
| RpS5a | FBgn0002590 | Yes | No | 1.058 | 0.262 | 37.4 | 102 |
| RpS6 | FBgn0261592 | Yes | Yes | -1.159 | 0.192 | 154 | 430 |
| RpS7 | FBgn0039757 | Yes | No | -1.011 | 0.273 | 217 | 595 |
| RpS8 | FBgn0039713 | Yes | No | 1.004 | 0.679 | 134 | 344 |
| RpS9 | FBgn0010408 | No | No | -1.579 | 0.013 | 229 | 375 |
| RpS10b | FBgn0285947 | Yes | No | -1.047 | 0.10 | 289 | 719 |
| RpS11 | FBgn0033699 | No | No | -1.599 | 0.0002 | 56.8 | 137 |
| RpS12 | FBgn0260441 | Yes | No | -1.007 | 0.891 | 121 | 214 |
| RpS13 | FBgn0010265 | Yes | No | 1.065 | 0.015 | 196 | 528 |
| RpS14a | FBgn0004403 | Yes | No | -1.112 | 0.236 | 581 | 1490 |
| RpS14b | FBgn0004404 | Yes | No | -1.366 | 0.038 | 191 | 535 |
| RpS15 | FBgn0034138 | Yes | No | -1.016 | 0.236 | 820 | 3040 |
| RpS15Aa | FBgn0010198 | Yes | Yes | -1.078 | 0.033 | 132 | 430 |
| RpS15Ab | FBgn0033555 | No | No | -1.078 | 0.033 | 32.1 | 64.8 |
| RpS16 | FBgn0034743 | Yes | No | -1.029 | 0.032 | 85.7 | 279 |
| RpS17 | FBgn0005533 | Yes | No | -1.050 | 0.184 | 167 | 622 |
| RpS18 | FBgn0010411 | Yes | No | -1.040 | 0.291 | 209 | 441 |
| RpS19a | FBgn0010412 | Yes | Yes | 1.011 | 0.323 | 196 | 610 |
| RpS20 | FBgn0019936 | No | No | -1.138 | 0.011 | 419 | 1260 |
| RpS21 | FBgn0015521 | Yes | No | 1.042 | 0.216 | 176 | 466 |
| RpS23 | FBgn0033912 | Yes | No | -1.198 | 0.0008 | 164 | 399 |
| RpS24 | FBgn0261596 | No | No | 1.028 | 0.272 | 240 | 690 |
| RpS25 | FBgn0086472 | Yes | No | -1.032 | 0.076 | 471 | 929 |
| RpS26 | FBgn0261597 | Yes | No | -1.043 | 0.021 | 268 | 717 |
| RpS27 | FBgn0039300 | Yes | Yes | -1.013 | 0.423 | 257 | 401 |
| RpS27A | FBgn0003942 | Yes | No | 1.018 | 0.355 | 235 | 528 |
| RpS28b | FBgn0030136 | Yes | No | -1.048 | 0.197 | 355 | 697 |
| RpS29 | FBgn0261599 | Yes | No | -1.079 | 0.091 | 328 | 909 |
| RpS30 | FBgn0038834 | Yes | No | -1.066 | 0.052 | 229 | 521 |
| Rack1 | FBgn0020618 | Yes | No | -1.091 | 0.003 | 435 | 1430 |
| Sta | FBgn0003517 | Yes | No | -1.068 | 0.008 | 198 | 596 |
| Vig2 | FBgn0046214 | Yes | No | 1.151 | 0.224 | 5.8 | 8.98 |
