## Supplementary material for "The Ribb-osome: Ribbon boosts ribosomal protein gene expression to coordinate organ form and function": Table S3

| **Gene** | **Forward Primer (5’to3’)** | **Reverse Primer (5’to3’)** | **Ann. Temp.** | **Amplicon** | **Source** |
| --- | --- | --- | --- | --- | --- |
| RpLP2 | ATCAAGGAGGGTCGCGAGAA | TAAGCGGTTGAATGTGGCGA | 55°C | 197 bp | This study |
| RpL11 | TCGGTATTCGCCGTAACGAG | CAACTTCGGTTTCGGCATCC | 55°C | 150 bp | This study |
| RpL19 | GGTCACTAAGCAGTCCGACC | TTTCCCTGGGAGCAGTCGTA | 55°C | 156 bp | This study |
| RpL28 | TGCTACACAACCAAGATCGCA | CCCCCGTCAAGGGAAAGAAG | 55°C | 209 bp | This study |
| RpS9 | TGGTCAACATCCCGTCGTTC | TAAGCAGTGGTAGCCAGCTG | 55°C | 181 bp | This study |
| RpS17 | GGTGCGTGGTATCTCCATCA | CAACAACTTTGGTCGTCGCA | 55°C | 197 bp | This study |
| RpS23 | CCTAACTCTGCCATCCGCAA | ATGCCGTCGGTGATATTCCC | 55°C | 165 bp | This study |
| Hsp70Ba | AGCGCACACTCTCCTCTAGC | GTCGTGGATCTGACCCTTGT | 55°C | 188 bp | Loganathan et al. |
| CG7130 | ACAAGATTCTGGGCATCGAG | CGCGCTTTTCCTTATCAAAG | 55°C | 168 bp | Loganathan et al. |
| CG33003 | ATATTTCCGAGGACCTTCAACTC | CCTCTGTGACAACTAACGGG | 55°C | 165 bp | Loganathan et al. |
| CLS | TGGAACTATTTGGATCACCCAG | CAGATATTTAGCGGAAATGCCC | 55°C | 154 bp | Loganathan et al. |
| Obp99b | TGCTTCGGTAGGTACTGTGT | AGGCCAGACCCAATAGGAGA | 55°C | 133 bp | Loganathan et al. |
| StnA | AAGACGAGGAGGACGACGAGTTC | TGGTTTTCCTGGTTCTTGGCCCG | 55°C | 176 bp | Loganathan et al. |
| Prosap | CTCCTCCTATTCCCGAACCT | GAACTACTCCCACTAGCCTG | 55°C | 156 bp | Loganathan et al. |
| Tlk | GCAGGAGTATTACGAGTACGA | GTTAAAGCGGGACTGATCCTC | 55°C | 162 bp | Loganathan et al. |

**Supplementary Table 3,** related to Figure 4.

A. Primers used for qRT-PCR validation of Rib targets.

B. Primers (5’to3’) used for generation of the *trh*-GAL4 line; related to Fig. S6

Forward*:* GGCCGCGGGCTACAATCCTTACAGTAACCTTAATCG

Reverse: GGGGATCCCTCGATATCTCAGTGTAAGAGGG
