## Supplementary material for "The Ribb-osome: Ribbon boosts ribosomal protein gene expression to coordinate organ form and function": Table S4

**Supplementary Table 4,** related to Figure 6.

Oligos used for EMSAs with 1) GST-Rib-PSQ DNA binding domain recombinant protein, 2) Rib-full length recombinant protein, 3) GST only and 4) dCrebA-full length recombinant protein.

| **Enhancer** | **Oligo** | **EMSAs with** |
| --- | --- | --- |
| RpL6 | CCCTGGTCACACCGCTCTAGACATAACCTTCCTTTTCCAATAATTTTTGT | 1, 2, 3 |
| RpL6-RC | ACAAAAATTATTGGAAAAGGAAGGTTATGTCTAGAGCGGTGTGACCAGGG | 1, 2, 3 |
| RpL7A | AACTGCCACTTCCTTTCTTTTTCGTTCCACGTTTCCGGTGAGTGCATGTG | 1, 2, 3 |
| RpL7A-RC | CACATGCACTCACCGGAAACGTGGAACGAAAAAGAAAGGAAGTGGCAGTT | 1, 2, 3 |
| RpL11 | CGCGGCCACACTATCCTTTCCTTCTTTTCTTACAAGTTTCCGGTGAGTGC | 1, 2, 3 |
| RpL11-RC | GCACTCACCGGAAACTTGTAAGAAAAGAAGGAAAGGATAGTGTGGCCGCG | 1, 2, 3 |
| RpL17 | ACTGCCATTTCCTTTCTTTTTCGTTCCACGTTTCCGGTGAGTGCATGTGC | 1, 2, 3 |
| RpL17-RC | GCACATGCACTCACCGGAAACGTGGAACGAAAAAGAAAGGAAATGGCAGT | 1, 2, 3 |
| RpL19 | TTCCGGCTGTCAAATTTTTCTTTCTTTCCTAGCCACGACGAGGTCGACAG | 1, 2, 3 |
| RpL19-RC | CTGTCGACCTCGTCGTGGCTAGGAAAGAAAGAAAAATTTGACAGCCGGAA | 1, 2, 3 |
| RpS3 | AACGTGTCTCGCGCGGGCACACTGGCCTCTTTCTTTTCTGCGCACCACGT | 1, 2, 3 |
| RpS3-RC | ACGTGGTGCGCAGAAAAGAAAGAGGCCAGTGTGCCCGCGCGAGACACGTT | 1, 2, 3 |
| RpS10b | ATCCCTTCCGTTTCCCTTTCTTTTCGTCTTCACGTTTCCGGTGAGTGCGT | 1, 2, 3 |
| RpS10b-RC | ACGCACTCACCGGAAACGTGAAGACGAAAAGAAAGGGAAACGGAAGGGAT | 1, 2, 3 |
| RpS17 | ACACAACCACCCTGGTCACACGGCATCCTTCTTTTTCTTTCGTTTCCGGC | 1, 2, 3 |
| RpS17-RC | GCCGGAAACGAAAGAAAAAGAAGGATGCCGTGTGACCAGGGTGGTTGTGT | 1, 2, 3 |
| RpS17* | ACACAACCACCCTGGTCACACGGCATCCGGCGGGGGCGGGCGGGGCCGGC | 1, 2, 3 |
| RpS17*-RC | GCCGGCCCCGCCCGCCCCCGCCGGATGCCGTGTGACCAGGGTGGTTGTGT | 1, 2, 3 |
| RpS18 | GGCAGCCACATTAATTCGGCCTTCTTTTCTTTTCCCGTTTCCGGTGAGTG | 1, 2, 3 |
| RpS18-RC | CACTCACCGGAAACGGGAAAAGAAAAGAAGGCCGAATTAATGTGGCTGCC | 1, 2, 3 |
| RpS27A | TTCAACTTCCTGTCACGCTGTGTGTGTGTTCTTTCTTTCCTTTGATTCTG | 1, 2, 3 |
| RpS27A-RC | CAGAATCAAAGGAAAGAAAGAACACACACACAGCGTGACAGGAAGTTGAA | 1, 2, 3 |
| P24.1 | CCGATAGCATATACGTGGGTCCGCAGTGC | 1, 2, 3, 4 |
| P24.1-RC | GCACTGCGGACCCACGTATATGCTATCGG | 1, 2, 3, 4 |
| TSN-2 | ATCTGATTTGGTGACGTTGGCACCCTGCA | 1, 2, 3, 4 |
| TSN-2-RC | TGCAGGGTGCCAACGTCACCAAATCAGAT | 1, 2, 3, 4 |

RC: Reverse Complement

RPS17*: Altered RPS17 sequence with G’s substituted for T’s in the promoter region
