## Supplementary material for "The Ribb-osome: Ribbon boosts ribosomal protein gene expression to coordinate organ form and function": Table S5

| GO term | Enrichment Score | PValue | Bonferroni | Benjamini | FDR |
| --- | --- | --- | --- | --- | --- |
| Transcription regulation | 5.08 | 1.90E-08 | 1.51E-05 | 5.32E-06 | 5.26E-06 |
| Calponin homology domain | 3.61 | 3.99E-06 | 0.001 | 0.001 | 0.001 |
| Zinc finger | 3.42 | 5.65E-05 | 0.016 | 0.001 | 0.001 |
| Pleckstin homology-like | 3.09 | 2.15E-06 | 0.003 | 0.003 | 0.003 |
| Kinase | 2.89 | 3.95E-06 | 0.01 | 1.25E-04 | 1.15E-04 |
| Src homology-3 domain | 2.79 | 2.73E-04 | 0.362 | 0.049 | 0.049 |
| Basic-leucine zipper domain | 2.59 | 9.33E-05 | 0.142 | 0.031 | 0.030 |
| Nuclear hormone receptor-type | 2.57 | 8.91E-05 | 0.091 | 0.016 | 0.016 |
| Transmembrane | 2.55 | 2.55E-04 | 0.070 | 0.005 | 0.005 |
| EF-hand-like domain | 2.11 | 9.71E-04 | 0.798 | 0.123 | 0.122 |

**Supplementary Table 5,** related to Supplemental Fig. 6

**DAVID analysis of genes bound by Ribbon in the embryonic Trachea – Top 10 annotation clusters**
